# Molecular Basis for Asynchronous Chain Elongation During Rifamycin Antibiotic Biosynthesis

**DOI:** 10.1101/2025.07.05.663307

**Authors:** Chengli Liu, Ryan C. West, Muyuan Chen, Whitaker Cohn, George Wang, Aryan M. Mandot, Selena Kim, Dillon P. Cogan

**Author notes:** Correspondence to Dillon P. Cogan., **Email:**. These authors contributed equally to this work (Chengli Liu, Ryan C. West). **Author Contributions:** C.L., R.C.W., W.C., G.W., A.M.M., S.K., and D.P.C. collected and analyzed experimental data. D.P.C. and M.C. produced cryo-EM maps and models. D.P.C., C.L., and R.C.W. wrote the initial manuscript which was revised by all authors.

## Abstract

The rifamycin synthetase (RIFS) from the bacterium *Amycolatopsis mediterranei* is a large (3.5 MDa) multienzyme system that catalyzes over 40 chemical reactions to generate a complex precursor to the antibiotic rifamycin B. It is considered a hybrid enzymatic assembly line and consists of an N-terminal nonribosomal peptide synthetase loading module followed by a decamodular polyketide synthase (PKS). While the biosynthetic functions are known for each enzymatic domain of RIFS, structural and biochemical analyses of this system from purified components are relatively scarce. Here, we examine the biosynthetic mechanism of RIFS through complementary crosslinking, kinetic, and structural analyses of its first PKS module (M1). Thiol-selective crosslinking of M1 provided a plausible molecular basis for previously observed conformational asymmetry with respect to ketosynthase (KS)–substrate carrier protein (CP) interactions during polyketide chain elongation. Our data suggest that C-terminal dimeric interfaces—which are ubiquitous in bacterial PKSs—force their adjacent CP domains to co-migrate between two equivalent KS active site chambers. Cryogenic electron microscopy analysis of M1 further supported this observation while uncovering its unique architecture. Single-turnover kinetic analysis of M1 indicated that although removal of C-terminal dimeric interfaces supported 2-fold greater KS-CP interactions, it did not increase the partial product occupancy of the homodimeric protein. Our findings cast light on molecular details of natural antibiotic biosynthesis that will aid in the design of artificial megasynth(et)ases with untold product structures and bioactivities.

**Significance Statement:** Bacteria use enzymatic assembly lines for the manufacture of complex and often medicinally active organic compounds. Their conserved modular design and biosynthetic logic suggest they evolved to be intrinsically reprogrammable. Yet, strategies to manipulate assembly-line product structures through protein engineering are still met with considerable challenges. This work investigates how a representative bacterial assembly line catalyzes an essential reaction during biosynthesis of the antibiotic rifamycin. Our data provide a molecular rationale for asynchronous C-C bond formation catalyzed by equivalent subunits of the homodimeric system while exposing new aspects of assembly line architecture. These findings represent a step closer towards the design of artificial assembly lines for sustainable production of user-defined chemicals.

## Introduction

In bacteria, specialized enzymatic assembly lines convert simple chemical building blocks from primary metabolism into structurally complex and medicinally important natural products(1, 2). Each assembly line consists of a group of multi-enzyme ‘modules’ that together coordinate stepwise maturation of biosynthetic intermediates across a defined sequence of active sites. The modular structure and behavior of these systems present a promising natural platform for directing biosynthesis of custom molecules via genetic reprogramming(3). Despite knowledge of hundreds of natural assembly lines that share this modular framework(4)—and bioinformatic inference of thousands more(5–7)—universal strategies for engineering new biosynthetic pathways while maintaining catalytic integrity remain limited. The recent discovery of a natural assembly line extending beyond 50 modules in length(8) combined with an approximate number of 10 canonical module functions(4, 9) implies that recombining these systems could theoretically encode biosynthetic pathways to >10^50^ unique products. Understanding the molecular mechanisms behind assembly-line biosynthesis will be key to approaching this enormous potential.

Here, we analyze the biosynthesis of a complex antibiotic precursor, proansamycin X (PAX), by the rifamycin synthetase (RIFS) in *Amycolatopsis mediterranei* as a model for understanding assembly-line mechanisms in bacteria (Fig. 1A)(10). RIFS is a nonribosomal peptide synthetase (NRPS)-polyketide synthase (PKS) hybrid encoded on five separate genes (*rifA–E*). The entire 3.5 MDa homodimeric assembly line contains an N-terminal NRPS loading module (LM) followed by ten PKS elongating modules (M1–M10) (Fig. 1B). The adenylation (A) domain of the LM initiates biosynthesis by selecting 3-amino-5-hydroxybenzoate (3,5-AHB) for ATP-dependent arylation of the 4′-phosphopantetheine (Ppant) cofactor of its adjacent substrate carrier protein (CP) domain. The 3-amino-5-hydroxybenzoyl-CP thioester product is then received by the ketosynthase (KS) domain of M1 through formation of a thioester linkage with its catalytic Cys residue, whereupon the intermediate initiates repeated elongation and modification by a decamodular PKS (Fig. 1C). This basic mechanism for incorporation of 3,5-AHB by an NRPS LM (consisting of an A-CP didomain), followed by transfer of the 3-amino-5-hydroxybenzoyl group onto the first KS domain of a multimodular PKS assembly line is shared across the structurally diverse family of ‘ansamycin’ natural products(11). In general, PKS modules from ansamycin assembly lines feature an embedded acyltransferase (AT) domain, making them ‘cis-AT’ type modules, as opposed to ‘trans-AT’ type modules which collaborate with separately encoded, or ‘standalone’, AT domains(6). The AT domain catalyzes transfer of an (α-substituted)malonyl group from an acyl-coenzyme A metabolite onto the Ppant thiol of its downstream CP domain, thereby supplying the nucleophilic substrate of KS-catalyzed Claisen condensation (elongation) with an electrophilic KS-bound thioester (see Fig. S1 for an illustration of the catalytic cycle of RIFS M1). Additional modifying domains, such as a ketoreductase (KR), dehydratase (DH), or enoylreductase (ER) may be present in the PKS module and usually catalyze NADPH-dependent reduction (KR/ER), dehydration (DH), or isomerization (KR/DH) of elongated polyketide intermediates before they are received by the downstream module’s KS. This process of elongation followed by modification continues at every active module in the assembly line until the final intermediate is covalently released through thioester cleavage by a cis-or trans-acting enzyme. Covalent release from RIFS occurs when the M10-bound linear intermediate undergoes unimolecular amidation between its aryl amine and thioester groups to afford PAX, the first macrolactam precursor of the antibiotic rifamycin B and several of its semisynthetic derivatives (Fig. 1A). This reaction is putatively catalyzed by RifF, an amide synthase homolog encoded directly downstream of RIFS (Fig. 1B)(12, 13). Genetic inactivation experiments predict that a second non-RIFS transformation occurs prior to formation of PAX involving aromatic hydroxylation of the M3-bound tetraketide intermediate by a putative flavin-dependent enzyme, Rif-Orf19 (Fig. 1B)(14). The hydroxyquinone product of this reaction sets the stage for oxidative cyclization into the bicyclic dihydronaphthoquinone ring observed in PAX and several post-M3 intermediates previously isolated(15).

**Figure 1.**
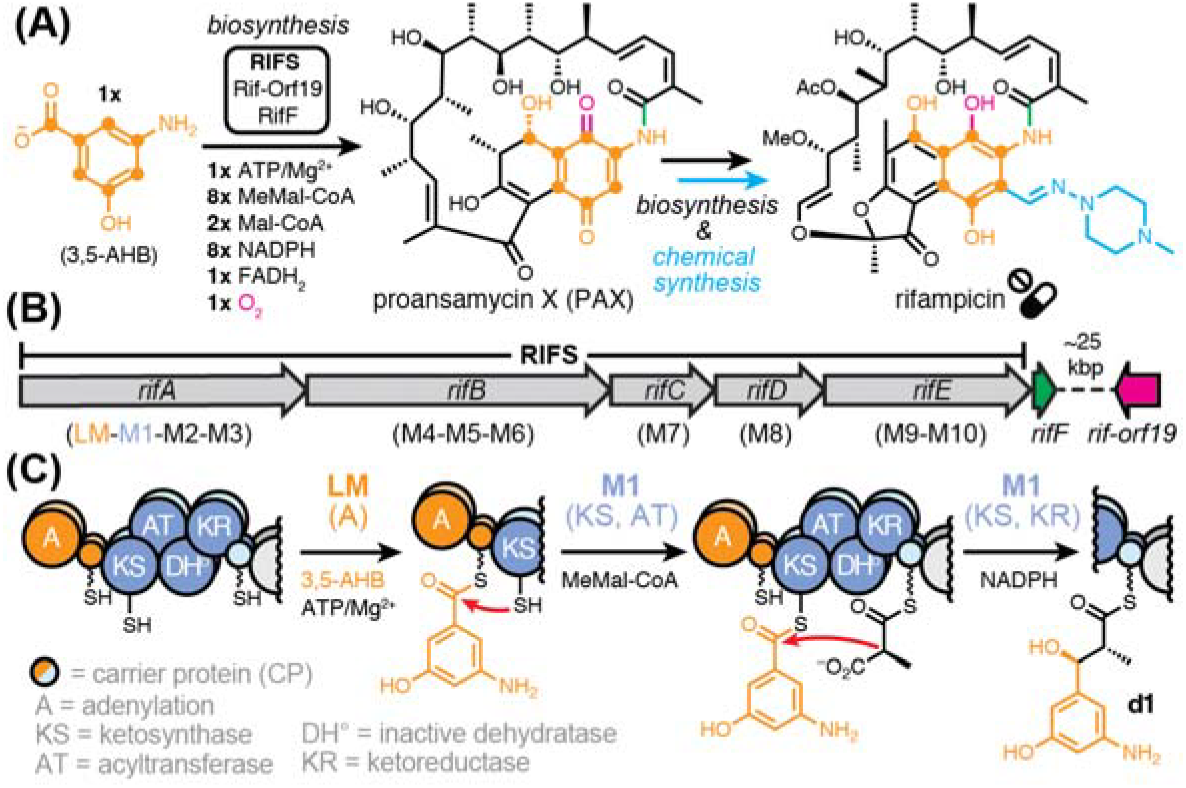
**(A)** Biosynthesis of rifamycin precursor, proansamycin X (PAX), by the RIFS assembly line in partnership with RifF and Rif-Orf19 (Mal = malonyl; MeMal = (2*S*)-methylmalonyl). The clinically-approved antibiotic rifampicin (a.k.a. rifampin) can be chemically synthesized from PAX-derived rifamycin B. **(B)** The genetic organization of RIFS (*rifA–E*), *rifF*, and *rif-orf19* in *Amycolatopsis mediterranei*. **(C)** Biosynthesis of diketide thioester (**d1**) en route to PAX by the LM and M1 of RIFS. The absolute stereochemistry of **d1** was inferred based on KR sequence motifs that predict stereochemical outcomes of β-ketoreduction and α-epimerization(48). Squiggly lines attached to sulfur atoms depict 4′-phosphopantetheine (Ppant) groups. The third transformation illustrates *inter*-molecular Claisen condensation between KS- and CP-tethered reactants from different subunits (distinguished by heavy and light shading).

Given the universality of 3,5-AHB incorporation into the ansamycin natural products and its involvement of a hybrid NRPS-PKS interface, we sought to investigate the mechanism of mixed aryl diketide thioester (**d1**) formation by RIFS (Fig. 1C). Admiraal et al. previously reconstituted **d1** formation *in vitro* from the RIFS LM-M1 bimodule while demonstrating that A-domain catalyzed arylation of the loading CP domain was rate-limiting compared with downstream reactions catalyzed by the PKS module, M1(16–18). Under single-turnover conditions, they observed that loading CP arylation proceeded to a roughly 3-fold higher extent than **d1** formation, implying that only one of the two PKS subunits of LM-M1 was fully active. Likewise, analysis of the erythromycin synthase determined that a homodimeric PKS module was roughly half-occupied with its growing polyketide intermediate under steady-state turnover conditions(19). These observations raise the questions of how and why equivalent pairs of PKS active sites are regulated such that only one is fully functional at a time. Recently reported structures of PKS modules in asymmetric states support the idea that catalysis in each subunit occurs asynchronously(20–22). Earlier studies also confirmed the potential for inter-subunit collaboration during KS-catalyzed chain elongation(23). Notwithstanding these advances, an accurate description of the sequence of steps and their associated subunits throughout a modular PKS’s catalytic cycle remains incompletely defined (Fig. S1).

We recently addressed how the KS domain engages its up- and down-stream CP partners by employing a thiol-reactive crosslinker to measure KS-CP interactions both within and between PKS modules of the erythromycin synthase(22). The reagent 1,3-dibromoacetone (DBA) was shown to react within seconds with native thiol groups of the homodimeric protein, thereby promoting site-selective crosslinking between the catalytic Cys and Ser-Ppant residues of the KS and CP domains, respectively (Fig. 2A). For a single PKS module, we expected that 0, 1, or 2 of its KS-CP pairs could become crosslinked, given that DBA should promote or prevent crosslinking based on inter-domain proximity and the fast observed rate of thiol alkylation(22). Biochemical and structural analyses indicated that no more than one KS-CP crosslink formed in each homodimeric module. Additional crosslinking experiments wherein CP domains were supplied as standalone proteins confirmed that this limit was only enforced when CP domains were covalently tethered to the PKS module (as they are in the natural system). These data, however, did not address the molecular basis for single CP occupancy, nor did they address the functional relevance of this phenomenon. Here, we provide a molecular explanation for asynchronous KS-CP interactions during intramodular polyketide elongation through the application of thiol-selective crosslinking of RIFS M1. Our data suggest that dimeric interfaces C-terminal to the CP domains impose a single KS-CP interaction limit per homodimeric module, as their removal promoted simultaneous crosslinking between both KS-CP pairs. Single-particle cryogenic electron microscopy (cryo-EM) analysis of M1 supported these conclusions while capturing snapshots of its unique module architecture. Finally, single-turnover kinetic analysis of M1 revealed, contrary to our expectation, that its C-terminal dimeric interface is not the sole determinant of partial product accumulation on the homodimeric protein. These findings establish a molecular basis for asymmetric interactions during the chain elongation stage of biosynthesis of a hybrid NRPS-PKS-derived natural antibiotic.

**Figure 2.**
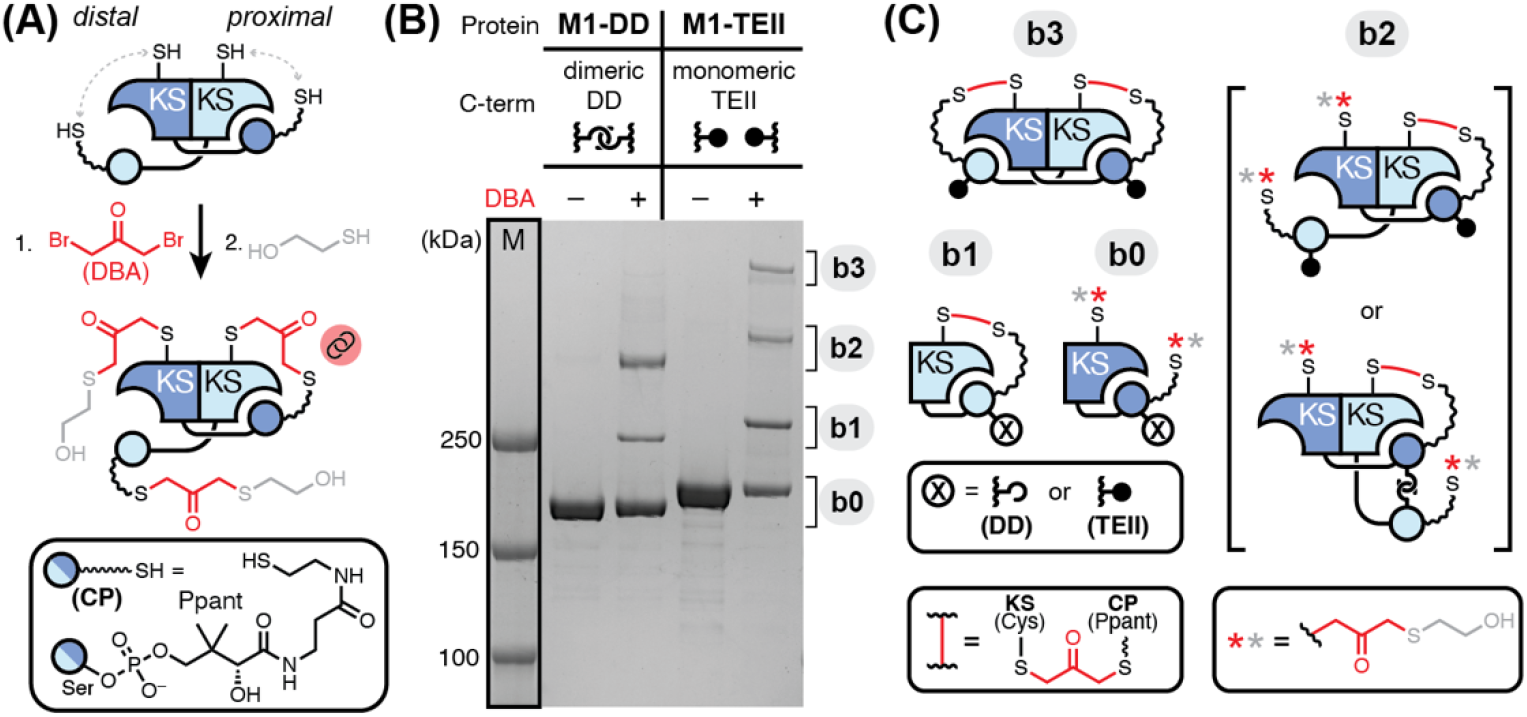
**(A)** DBA addition to a homodimeric *holo*-form PKS module (step 1) followed by quenching with excess thiol (step 2) permits proximity-dependent crosslinking between thiols present on KS and CP domains. For simplicity, only one possible crosslinking outcome is shown, and domains not involved in crosslinking have been omitted (Ppant = 4′-phosphopantetheine). **(B)** SDS-PAGE analysis following addition of DBA to M1-DD and M1-TEII, which are derivatives of the first PKS module of RIFS, revealed the appearance of low-mobility crosslinked bands (**b1**–**b3**) relative to un-crosslinked material (**b0**). M1-DD contains a C-terminal dimeric docking domain (DD), whereas M1-TEII contains a C-terminal monomeric type II thioesterase (TEII) ‘RifR’ associated with the rifamycin biosynthetic gene cluster(34). **(C)** Cartoon structures of **b0**–**b3** illustrate their *intra*- or *inter*-molecular crosslinked or un-crosslinked nature.

## Results

To address whether the observed single KS-CP interaction limit(22) applies to PKS modules from two phylogenetically distinct assembly lines (i.e., the erythromycin and rifamycin synth(et)ases), we isolated C-terminally His_6_-tagged RIFS M1 for DBA crosslinking analysis. The recombinant protein (M1-DD) was also tagged with short α-helical N-/C-terminal dimeric docking domains (DDs) from the erythromycin synthase to enable antibody complexation and inter-modular communication, as before(24, 25). Co-expression of M1-DD with Sfp(26, 27) in *E. coli* BAP1(28) allowed for post-translational 4′-phosphopantetheinylation at Ser1538 of its CP domain from cellular coenzyme A (i.e., conversion from its *apo-* to *holo*-form). Addition of DBA to *holo*-M1-DD followed by SDS-PAGE analysis revealed partial conversion of the starting material into two products, bands 1 (**b1**) and 2 (**b2**), with reduced mobility relative to the uncrosslinked material (**b0**) (Fig. 2B). Similarity of **b1** and **b2** to previously crosslinked modules from the erythromycin synthase(22) suggested that *intra*- and *inter*-molecular KS-CP crosslinks had occurred, respectively (Fig. 2C). To confirm the site-selectivity of DBA crosslinking, we prepared *apo*-M1-DD by expressing the same protein in *E. coli* BL21(DE3) (which lacks Sfp) and variants of *holo*-M1-DD whose catalytic KS Cys (C203) residue had been substituted with Ala or Ser. Crosslinking analysis of these M1-DD variants alongside wild-type *holo*-M1-DD revealed that thiols of the Ppant and catalytic Cys from the CP and KS domains, respectively, were principally involved in formation of **b1** and **b2** (Fig. S2). Although minor in comparison, the appearance of crosslinked bands similar to **b1** and **b2** in the *holo*-M1-DD(C203(A/S)) variants implied that other nucleophilic residues near the KS active site can participate in DBA crosslinking (Fig. S2). The observation that RIFS M1-DD could be chemically converted into the above crosslinked products also indicated that its catalytically inactive dehydratase domain (DH°), not present in previously tested PKS modules(22), was not influential in enforcing the single KS-CP crosslinking limit. Thus, asymmetry with respect to intramodular KS-CP interactions appears to be a common feature of these two distinct PKS assembly lines and module architectures.

To understand the basis for this phenomenon, we considered a previous model proposed by Bagde et al. wherein C-terminal dimeric interfaces serve to link CP domains non-covalently and therefore preclude their simultaneous engagement with spatially separated KS active sites(30). This model is so far in agreement with available cryo-EM structures of homodimeric PKS modules featuring intramodular KS-CP interactions and a C-terminal dimeric domain(22, 30–32). Hence, we predicted that replacement of a PKS module’s C-terminal dimeric domain with a monomeric domain might allow CP domains to interact and crosslink with their KS partners simultaneously. Accordingly, we substituted the C-terminal DD of RIFS M1-DD, which is dimeric(33), with the type II thioesterase (TEII) associated with the rifamycin biosynthetic gene cluster (a.k.a. RifR), which is monomeric(34); thereby affording M1-TEII (Fig. 2B). Similar isolation of *holo*-form M1-TEII followed by DBA addition and SDS-PAGE analysis again revealed low-mobility bands **b1** and **b2** (with corresponding mass shifts from the larger TEII domain) in addition to a newly observed band (**b3**) with even further reduced mobility (Fig. 2B–C). To investigate whether **b3** was formed through a non-specific reaction with the TEII, we prepared a version of M1-DD featuring a 38-residue deletion corresponding to its dimeric α-helical motif of the C-terminal DD (M1-ΔDD). DBA addition to M1-ΔDD produced a new band relative to crosslinked M1-DD that migrated similarly to **b3** in M1-TEII, supporting the idea that DD was inhibitory to the formation of this species (Fig. S3). A distribution of low-mobility bands resembling **b1**–**b3** was previously observed following KS-CP crosslinking of the animal fatty acid synthase (FAS) which possesses an overall homodimeric structure and domain architecture like that of bacterial PKSs(35). Notably, the C-terminal thioesterase (TE) domain of animal FAS by itself is monomeric(36). The reported structural assignments of the **b1** and **b2** equivalents in crosslinked animal FAS are similar to those represented in Figure 2C and characterized earlier(22). Consistent with the model from Bagde et al.(30), the equivalent of **b3** in crosslinked animal FAS was found to be a crosslinked homodimer bearing two inter-molecular KS-CP crosslinks, suggesting similar species were formed after crosslinking M1-TEII and M1-ΔDD (Figs. 2 and S3).

To gain structural support of these putative crosslinked states, we applied our previous methodology for analyzing PKS modules by single-particle cryo-EM(22, 31). The M1-DD protein harbored an N-terminal α-coiled-coil epitope of an antibody fragment (F_ab_) ‘1B2’ previously shown to enhance the particle quality of homodimeric PKS modules(30, 31, 37). The module-F_ab_ complex (M1-DD-1B2) was therefore isolated (Fig. S4) and subjected to cryo-EM analysis, which resulted in a 2.86 Å consensus map from 1.3 million particles; refined without symmetry (Fig. S5).

This cryo-EM map featured prominent density for two F_ab_ heterodimers bound to the homodimeric KS-AT core of M1-DD. To assess sample heterogeneity, we subjected the consensus map and particles to 3D classification(38). A majority of the output 3D class averages displayed a region of density near the C-terminus of the KS-AT core that resembled previously reported crystal structures of dimeric DHs(39, 40) (Fig. S5). Further map refinement and model building of the M1-DD-1B2 complex supported assignment of this region as the dimeric DH°, representing the first structure of a dehydratase dimer embedded between an AT and KR domain within a PKS module. Two noteworthy cryo-EM maps of M1-DD-1B2 emerged from 3D classification and subsequent homogenous refinement (Fig. 3A–D). One of them features a single AT-CP interaction (3.96 Å; 37,950 particles), consistent with its conformation during AT-catalyzed extender unit transacylation (*transacylation-mode*; Figs. 3A–B and S5–S7). The other one features a single KS-CP interaction (3.22 Å; 177,509 particles), consistent with its conformation during KS-catalyzed polyketide chain elongation (*elongation-mode*; Figs. 3C–D, S5, and S8). In both states, map and model fitting guided by AlphaFold 3 supported the existence of electrostatic and hydrophobic interactions nearby and within the KS active site tunnel occupied by the Ppant cofactor (Fig. 3B and 3D)(41). Importantly, no more than one KS- or AT-bound CP was observed in all 3D class averages of M1-DD-1B2 (Fig. S5). Comparison of the *transacylation-mode* (Fig. 3A–B) and *elongation-mode* (Fig. 3C–D) structures suggested that their interconversion would be met with a 30° (or 150°) rotation of the DH° dimer (or KS-AT core) relative to the module’s pseudo-C2 axis of symmetry (Fig. 3E). Assuming that KS-catalyzed elongation occurs between substrates bound to different subunits—in accordance with previous findings (Fig. 1C)(23, 31)— the simplest interpretation is that the CP domain interacts *intra*-molecularly with its partner AT to receive a methylmalonyl extender unit before transiting to the nearest KS active site for *inter*-molecular elongation (Fig. 3E). Conversely, extender unit transacylation may occur *inter*-molecularly followed by *intra*-molecular elongation, or both reactions may occur on one or the opposite subunit (i.e., both are *intra-* or *inter*-molecular). Further experiments will be required to differentiate among these four possible AT-to-KS CP trajectories. Analysis of the *elongation-mode* cryo-EM map supported the existence of both *intra-* and *inter*-molecular KS-CP interactions, corresponding to **b1** and **b2**, respectively (Fig. S9).

**Figure 3.**
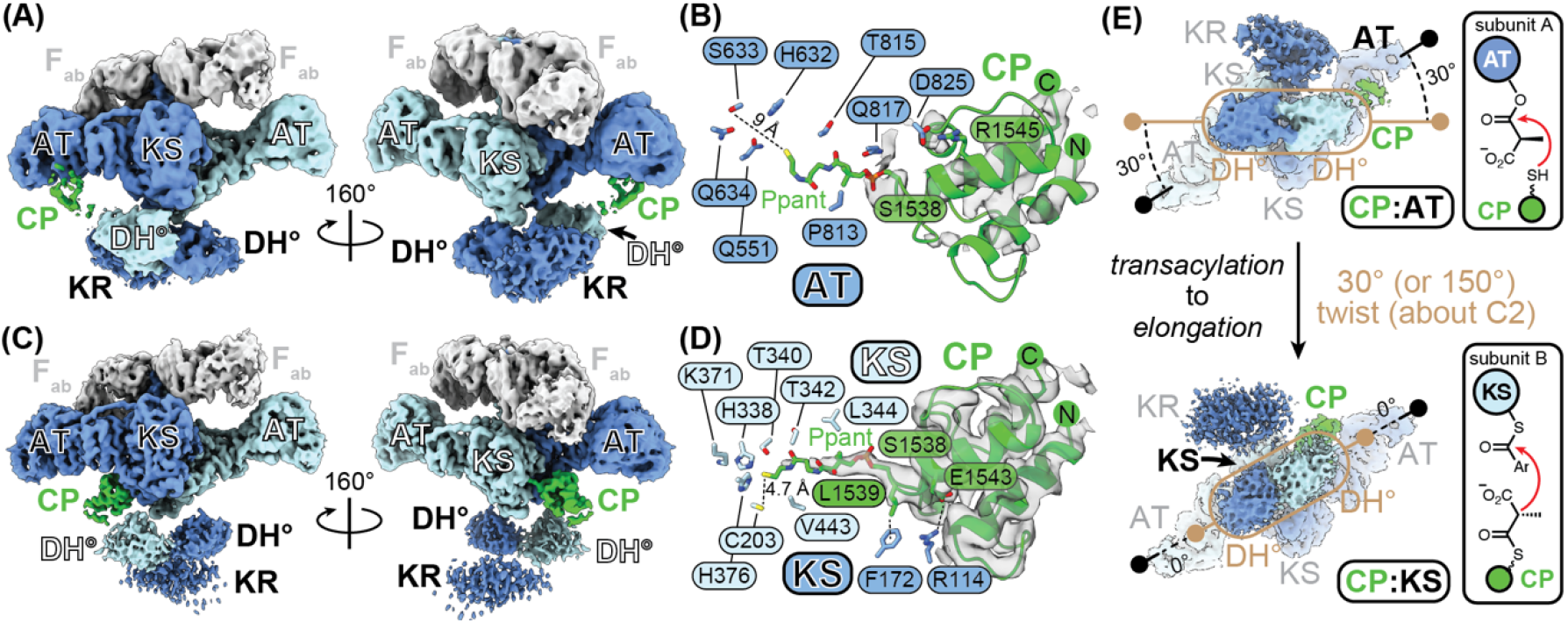
**(A)** The overall cryo-EM map of RIFS M1-DD bound to F_ab_ 1B2 (M1-DD-1B2) in the *transacylation-mode* and **(B)** modeled interactions between its 4′-phosphopantetheine (Ppant)-attached CP and AT domains at 3.96 Å gold-standard Fourier shell correlation (GSFSC) resolution (Figs. S5 and S6). Focused refinement produced a supplemental cryo-EM map with more prominent density for the CP domain in the *transacylation-mode* structure (Fig. S7). **(C)** The overall cryo-EM map of M1-DD-1B2 in the *elongation-mode* and **(D)** modeled interactions between its Ppant-attached CP and KS domains at 3.22 Å GSFSC resolution (Figs. S5 and S8). Side chain coordinates shown in panels B and D were approximated by AlphaFold 3, as the local resolution was insufficient for experimental map-based modeling (Figs. S6 and S8)(41). **(E)** Comparison of the *transacylation-* and *elongation-mode* structures of M1-DD-1B2 highlights potential motion during the catalytic cycle involving a 30° (or 150°) twist of the DH° dimer (or KS-AT dimer) about the pseudo-C2 axis of module symmetry.

To confirm or refute the putative **b3** structure harboring two *inter*-molecular KS-CP interactions (Fig. 2B–C), we prepared DBA-crosslinked M1-TEII bound to 1B2 (CL-M1-TEII-1B2) for similar cryo-EM analysis. Size-exclusion chromatography and SDS-PAGE supported the existence of a crosslinked module-F_ab_ complex featuring **b1**–**b3** (Fig. S4). Roughly 45% of the total particles of CL-M1-TEII-1B2 produced a cryo-EM map (3.96 Å; 91,575 particles) consisting of unambiguous density for both CP domains bound with their KS partners (Figs. 4A and S10–S11). DBA-dependent formation of **b3** in M1-TEII (but not in M1-DD) was therefore attributed to its absence of a C-terminal dimeric interface which enabled 2-fold KS-CP interactions (Fig. 2B–C). A similar 3D class average contained density for only one KS-bound CP domain, supporting the existence of **b1** and **b2** in the crosslinked sample mixture (Figs. S4 and S10). The cryo-EM map of CL-M1-TEII-1B2 also highlighted an unexpected interaction (supported by AlphaFold 3) between the unstructured KR-CP linker and a shallow groove of the opposite subunit’s DH° domain (Figs. 4B and S12). Therein, putative electrostatic contacts occur between Arg1480/Arg1481 of the KR-CP linker and Asp938/Asp1078 of the DH°, respectively. Notably, three out of four of these residues are highly conserved in PKS modules with similar domain architectures (Figs. 4C and S13). Overall, cryo-EM analysis of CL-M1-TEII-1B2, which featured 2-fold (pseudo-C2 symmetric) KS-CP interactions, provided additional support of the idea that dimeric interfaces at the C-termini of PKS modules act to confine both CP domains to a single KS active site cleft at a time. Moreover, this phenomenon appears to be independent of the type of dimeric interface employed, as multiple C-terminal domains have been associated with single KS-CP occupancy: namely, (i) the ‘post-AT dimerization element’ present in Lsd14(30), (ii) the type I TE of erythromycin synthase module 6(22, 31, 42), and (iii) the α-helical DD of erythromycin synthase module 2(22, 31, 33). A recent publication also highlighted single and double KS occupancy by CP domains in PKS modules from the colibactin-producing assembly line. Importantly, the observed 2-fold KS-CP interactions correlated with the absence of a C-terminal dimeric interface(32).

**Figure 4.**
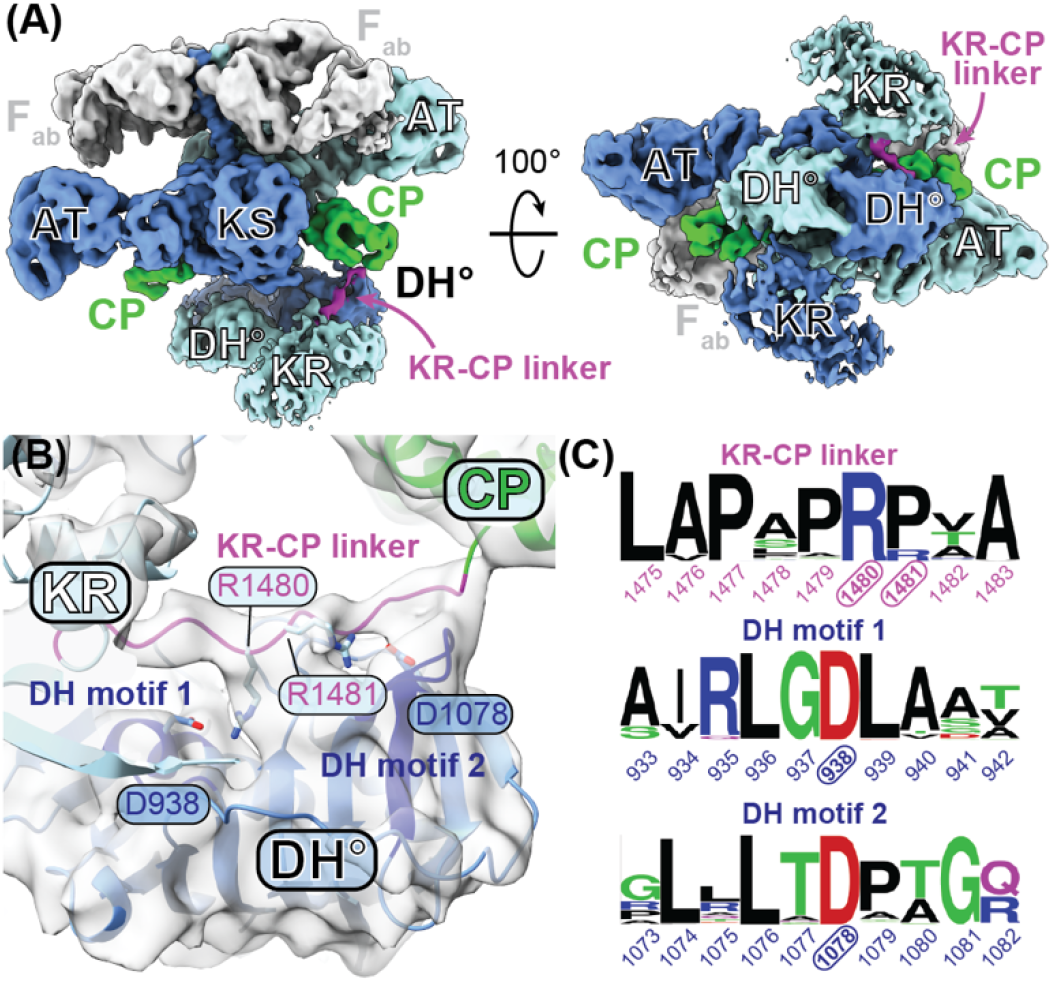
**(A)** The overall cryo-EM map of crosslinked RIFS M1-TEII bound to F_ab_ 1B2 (CL-M1-TEII-1B2) at 3.96 Å GSFSC resolution (Figs. S10–S11). A notable region of density corresponding to the KR-CP linker is highlighted in purple. Focused refinement produced a supplemental cryo-EM map with slightly improved resolution of the KR-CP linker (Fig. S12). The equivalent linker in the opposite subunit was not well resolved in either map. **(B)** Two Arg residues of the KR-CP linker (R1480 and R1481) were predicted by AlphaFold 3 to make electrostatic contacts with D938 and D1078 of the DH° domain from the opposite subunit. **(C)** WebLogos displaying residue type and frequency at regions containing these residues (i.e., KR-CP linker and DH motifs 1 & 2) were generated from a multiple sequence alignment of 250 homologs of the DH°-KR-CP fragment of RIFS M1 (WP_013222547.1)(49). From this analysis, three out of four of the residues putatively involved in electrostatics are invariant, whereas one of them (R1481) is often replaced by a Pro (see Fig. S13 for a complementary ConSurf analysis of sequence conservation mapped onto the DH°-KR-CP structure)(50).

Having established a plausible basis for single KS occupancy by a downstream CP domain in a homodimeric PKS module, we questioned the function of this behavior in the context of the PKS’s catalytic cycle. We considered that the C-terminal dimeric interface of M1-DD might prevent it from utilizing both subunits concurrently. From this, we hypothesized that the presence of a C-terminal dimeric interface would only allow complete polyketide processing in one catalytic subunit, whereas its absence would permit complete processing in both. If true, then M1-TEII would be expected to produce two aryl diketide thioester (**d1**) products following substrate incubation (sans product release from the module), whereas M1-DD would only produce one (Fig. 1C). As expressed above, evidence for partial module occupancy (i.e., ≤1 product per 2 subunits) has been observed in PKS modules from both the rifamycin(17) and erythromycin(19) synth(et)ase assembly lines. Notably, each of them harbored a C-terminal dimeric domain.

To test this prediction, we sought to develop a single-turnover kinetic assay capable of measuring the module-bound aryl diketide thioester (**d1**). Admiraal et al. previously demonstrated that, with addition of ATP/Mg^2+^, methylmalonyl coenzyme A (MeMal-CoA), and NADPH substrates, the RIFS LM-M1 bimodule could convert benzoate, a simple analog of its native starter unit, into 3-hydroxy-2-methyl-3-phenylpropanoyl-CP thioester (**d2**), the corresponding diketide analog of **d1** (Figs. 1C and 5A)(17). Hence, we obtained an authentic standard of the hydrolyzed product of **d2** (**d2**′) to enable its quantitation from hydrolysates of enzymatic reactions by liquid chromatography-tandem mass spectrometry (LC-MS/MS). We then purified recombinant forms of the RIFS LM-M1 bimodule C-terminally fused with DD or TEII (LM-M1-DD or LM-M1-TEII*) to measure the influence of C-terminal domains on enzymatic **d2** formation. In these experiments, we employed an inactive variant of the TEII domain—created by replacing its active site Ser with Ala(34) (denoted TEII*)—to prevent potential enzymatic release of **d2** and therefore ensure that the protein was limited to a single catalytic cycle per active subunit. To minimize possible steric effects of TEII* on enzymatic activity, a longer (G_4_S)_8_ linker between the CP and TEII* domains relative to M1-TEII was also employed in LM-M1-TEII*. DBA crosslinking reactions indicated that these changes remained compatible with **b1**–**b3** production and thus 2-fold KS-CP interactions (Fig. S3). Enzymatic production of **d2** was initiated by combining *holo*-form LM-M1-DD or LM-M1-TEII* with benzoate, ATP/Mg^2+^, MeMal-CoA (racemate), and NADPH and quenched at various time points from 1 min to 2 h by addition of KOH and heating (Fig. 5A). LC-MS/MS quantification of **d2**′ over time confirmed that both LM-M1-DD and LM-M1-TEII* were catalytically active, whereas LM-M1-DD(C802A), which lacked the essential Cys of the KS domain, exhibited no detectable activity (Fig. 5B). The maximum concentration of **d2′**′ approached roughly 30% of the concentration of protein in the LM-M1-DD containing reaction (Figs. 5B and S14), recapitulating the previous observation that protein-bound diketide (**d2**) accumulated sub-stoichiometrically prior to hydrolysis (despite presence of a large excess of substrates)(17). Interestingly, a nearly identical maximum diketide occupancy was observed in reactions containing LM-M1-TEII* (Figs. 5B and S14). Thus, contrary to our expectation, the extent of **d2** formation did not increase when the C-terminal dimeric domain of the PKS module was replaced with a monomeric domain. This implied that the C-terminal dimeric interface of LM-M1-DD was not the sole determinant of incomplete polyketide processing by the homodimeric protein.

**Figure 5.**
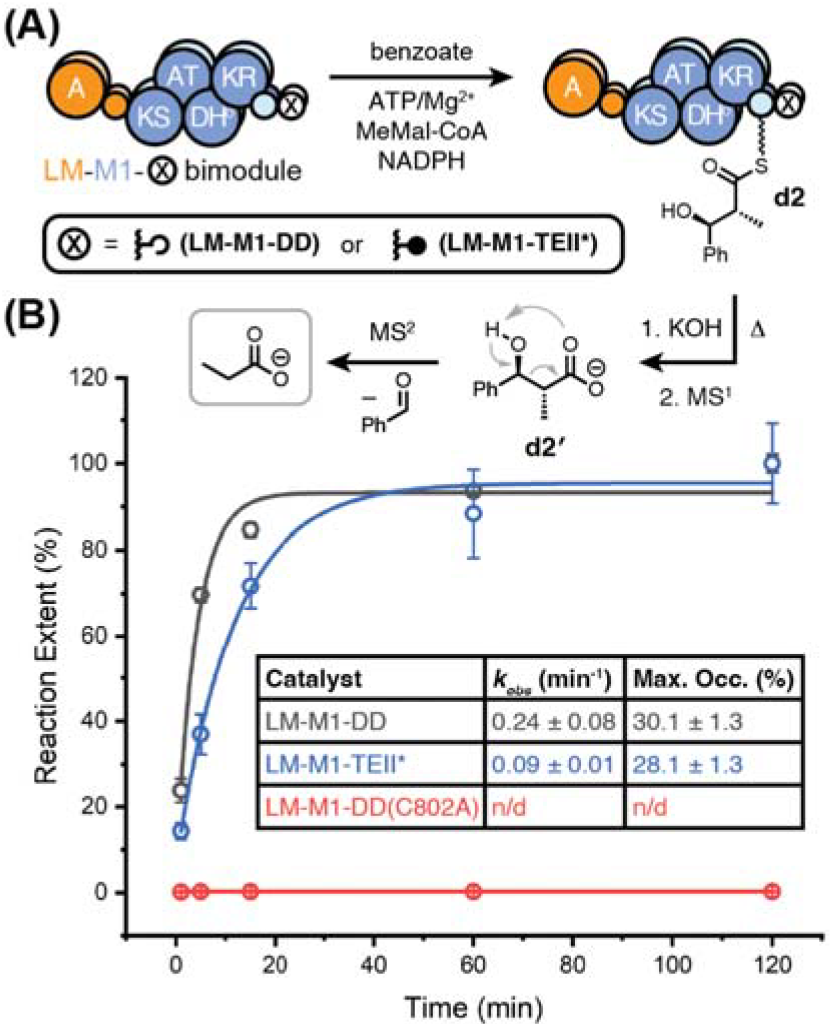
**(A)** Enzymatic production of covalently anchored diketide (**d2**) by the RIFS LM-M1-X bimodule, where X implies a C-terminal DD or TEII*. KOH and heat treatment of the enzyme-bound product liberated **d2**′ which was detected by LC-MS/MS. The primary fragment ion of **d2**′ (*m/z* = 73, presumed to be propionate; boxed in gray) was harnessed for quantification along with a secondary fragment ion (*m/z* 117, Fig. S14). **(B)** Normalized reaction extents are plotted for LM-M1-DD, LM-M1-TEII*, and LM-M1-DD(C802A) based on fragment ion counts over time. The data were fit to a single-phase exponential growth function to obtain approximate rate constants (*k*_*obs*_). LC-MS/MS quantification of **d2**′ standards in the presence of LM-M1-DD or LM-M1-TEII* was applied to infer the maximum percent occupancy of each protein with **d2** in enzymatic reactions (Fig. S14).

The observation that LM-M1-DD catalyzed **d2** formation approximately 2-to 3-fold faster than LM-M1-TEII* was also unexpected, considering that both proteins harbored identical LM-M1 fragments and were assumed to be equally rate-limited by A-domain benzoylation(16–18). We therefore isolated the rates of M1-catalyzed **d2** formation by pre-incubating LM-M1-DD or LM-M1-TEII* with benzoate and ATP/Mg^2+^ (to build up equal concentrations of the benzoylated protein) prior to addition of MeMal-CoA and NADPH co-substrates. Similar kinetic analysis of **d2**′ over time again revealed that LM-M1-DD outperformed LM-M1-TEII* with an approximate 2-fold greater rate constant (Fig. S15), suggesting that TEII* and/or its associated (G_4_S)_8_ linker were inhibitory to M1-catalyzed **d2** formation. Together, our data indicate that LM-M1-DD catalyzed **d2** formation faster than LM-M1-TEII* but up to the same fractional occupancy after a single catalytic cycle, despite the ability of the TEII* domain to confer 2-fold higher KS-CP interactions in the PKS module (Fig. S3).

## Discussion

Our analysis of the RIFS biosynthetic assembly line provides a molecular basis for previously observed conformational asymmetry in homodimeric PKS modules during KS-catalyzed C-C bond formation(30–32). In each of these cited cases, PKS modules bearing C-terminal domains with dimeric interfaces were employed. We showed with protein crosslinking and single-particle cryo-EM that simple replacement of a C-terminal dimeric α-helical DD with a monomeric TEII domain in RIFS M1 promoted a 2-fold increase in KS-CP interactions (Fig. 2). This observation mirrored previous analysis of an animal FAS, wherein similar 2-fold KS-CP interactions were observed for the PKS-like specimen terminating in a monomeric TE domain(35). Our cryo-EM analysis of RIFS M1 also provided previously unseen snapshots of a PKS module containing a KS-AT-DH°-KR-CP domain architecture. The characteristic shape of the DH° dimer was readily observed in cryo-EM maps of RIFS M1, which in turn provided a fortuitous tracer for measuring a putative structural transition during the catalytic cycle (Fig. 3). Our single-turnover kinetic analysis of the RIFS LM-M1 bimodule argued that even when both CP domains can engage their KS partners simultaneously—by virtue of attachment with a C-terminal monomeric TEII—only one of them facilitates the complete set of reactions leading to the diketide product (**d2**) (Fig. 5). We emphasize that our results did not rule out the possibility that C-terminal monomeric domains could allow two rounds of elongation per catalytic cycle with only one round of KR-catalyzed ketoreduction, as our detection method did not account for the pre-mature β-ketoacyl thioester analog of **d2**. In any case, our data suggest that other factors beyond the migrational independence of CPs control the limited extent of polyketide processing either during or following polyketide chain elongation. One possibility is that KS-catalyzed elongation in one catalytic subunit allosterically disengages the activities of its adjacent subunit, regardless of any effects of C-terminal domains on CP mobility. Such a scenario hearkens back to the previously discovered ‘turnstile’ mechanism, wherein KS-catalyzed elongation is coupled with temporary inhibition of KS activity(19) (Fig. S1). Despite multiple lines of evidence in favor of this mechanistic model, it remains unverified whether one or two rounds of elongation are required to achieve the fully KS-inhibited, or ‘turnstile closed’, conformation(31). While theoretically introducing TEII* into the C-terminus of LM-M1 could have increased the extent and/or rate of **d2** formation—given its promotion of 2-fold higher KS-CP interactions—we observed no effect on the extent and rather a reduced rate, relative to reactions with LM-M1 terminating in a more natural dimeric DD (Fig. 5). This finding may reflect an evolved preference of multi-functional PKS modules to utilize one subunit at a time for increased fidelity, unlike their mono-functional animal FAS counterparts which employ both subunits simultaneously to achieve their extraordinarily fast rates of catalysis(35, 43–47). Overall, this work deepens our understanding of molecular factors that control asynchronous C-C bond formation during modular polyketide antibiotic biosynthesis in bacteria.

## Acknowledgments

The authors thank Htet Khant and Tomasz (Tomek) Osinski at the University of Southern California for their assistance with cryogenic electron microscopy data acquisition and processing, respectively. Funding was provided to D.P.C. by a USC Mann School of Pharmacy Intramural Research Award and to M.C. by the National Institutes of Health (5R01GM150905-02).

## Data Availability and Sharing Plan

Associated atomic coordinates and cryo-EM maps have been deposited to the Protein Data Bank under accession codes 9PAT (*transacylation-mode*), 9PAV (*elongation-mode*), and 9PC6 (*CL-M1-TEII-1B2*) and to the Electron Microscopy Data Bank under accession codes EMD-71445, EMD-71446, and EMD-71497, respectively. All materials used in this study that are not commercially available can be made available by the authors upon request.

