## Supporting Information for "Molecular Basis for Asynchronous Chain Elongation During Rifamycin Antibiotic Biosynthesis"

Dillon P. Cogan

**This PDF file includes:**

Materials and Methods

Figures S1 to S15

Tables S1 to S4

Protein Sequences

SI References

**Materials and Methods**

**Materials.** Bacterial growth media, enzymes, purification resins, Amicon Ultra Centrifugal Filters, and other chemicals were purchased from Thermo Fisher Scientific, Avantor (VWR International), or Millipore Sigma. Antibiotics, isopropyl β-D-1-thiogalactopyranoside (IPTG), dithiothreitol (DTT), and tris-(carboxyethyl) phosphine (TCEP) were purchased from Gold Biotechnology. (2*R*,3*S*)-3-hydroxy-2-methyl-3-phenylpropanoic acid (**d2′**, Fig. 5) was purchased from Enamine (EN300-8130951). Cryogenic electron microscopy (cryo-EM) grids were purchased from Electron Microscopy Sciences or MiTeGen. Polyacrylamide gels were purchased from Thermo Fisher Scientific or Bio-Rad. DNA oligonucleotides were purchased from Integrated DNA Technologies, and DNA sequencing services were provided by Azenta Life Sciences. Microbial strains used in this work were provided by the USDA-ARS Culture Collection (NRRL).

**General Methods.** Protein purification was carried out using an ÄKTA Pure chromatography system (Cytiva Life Sciences). Unless otherwise stated, protein concentrations were determined using the Bradford assay with bovine serum albumin standards by measuring absorbance at 595 nm in 1 cm cuvettes using a NanoDrop OneC (Thermo Fisher Scientific)(1). DNA concentration and purity were assessed by using the stage of the NanoDrop OneC (Thermo Fisher Scientific) and measuring absorbance at 260/280 nm. Gels were imaged on a Bio-Rad ChemiDoc Imaging system.

**Genomic DNA Extraction.** *Amycolatopsis mediterranei* NRRL B-3240 obtained from NRRL as a freeze-dried pellet was streaked onto a plate of ISP Medium No. 4 (HiMedia) and grown at 28 °C for four days. The resulting colonies were scraped from the surface of the plate, and genomic DNA (gDNA) therein was extracted using the E.Z.N.A. Bacterial DNA Kit (Omega Bio-tek) according to the manufacturer’s protocol.

**Production of Plasmids for Protein Overexpression.** DNA oligonucleotides (0.3 µM) and template (4 ng/µL: gDNA; 0.4 ng/µL: plasmid DNA) were used to amplify the protein-coding regions with CloneAmp HiFi PCR Premix (Takara Bio USA) supplemented with 1 M betaine and 3% (ν/ν) dimethyl sulfoxide (DMSO) by employing a touchdown PCR protocol(2). Amplified products were analyzed and purified by agarose gel electrophoresis and ligated with similarly prepared pET21- or pET28-derived DNA fragments via In-Fusion Cloning (Takara Bio USA) to create circular DNA plasmids (Table S1). Initial plasmid products were used to transform *Escherichia coli* Stellar™ competent cells (Takara Bio USA), and the resulting transformants were grown for 12 h at 37 °C (220 rpm) in 5–10 mL LB (Miller) broth supplemented with 100 µg/mL carbenicillin. Plasmids were isolated from the liquid cultures via EasyPrep™ (Bioland Scientific LLC) and verified by Sangar sequencing prior to downstream use.

**Protein Expression and Purification.** Purification of the F_ab_ 1B2 was performed as before(3). For purification of all other proteins, expression plasmids (Table S1) were used to transform *E. coli* BL21(DE3) or BAP1(4) competent cells to generate proteins in their *apo* or *holo* forms, respectively. Single colony transformants were selected to inoculate 10–15 mL LB (Miller) broth supplemented with appropriate antibiotic (100 µg/mL carbenicillin or 50 µg/mL kanamycin) and grown for 14 h at 37 °C (220 rpm). Two mL of the resulting cell suspension were used to inoculate every 1 L of similar LB + antibiotic medium and grown at 37 °C (220 rpm) until an optical density at 600 nm (OD_600_) of ≈0.5 was achieved (3–6 L total). Cell cultures were cooled for 15 min in an ice bath before adding 0.25 mM IPTG and continuing growth at 18 °C (220 rpm) for 18–20 h. Cells were harvested by centrifugation at 3,000 × g for 30 min and resuspended in 5 mL of 0.45 M NaCl, 10 mM imidazole, 50 mM NaH_2_PO_4_, 20% glycerol, pH 7.8 (NaOH) per liter of cell culture. Cells were lysed by sonication with a Branson Sonifier 450, and lysates were clarified by centrifugation at 23,400 × g in a Sorvall RC 5B centrifuge. Clarified lysates were added to HisPur Ni-NTA resin (Thermo Fisher Scientific) in a glass Econo-Column (Bio-Rad) equilibrated with 0.3 M NaCl, 50 mM imidazole, 50 mM NaH_2_PO_4_, 10% glycerol, pH 7.8 (NaOH) (wash buffer) at a ratio of 1 mL resin per liter of cell culture. The protein-bound resin was washed with 100 mL of wash buffer in 4 × 25 mL increments before adding 50 mL of 40 mM NaCl, 500 mM imidazole, 50 mM NaH_2_PO_4_, 10% glycerol, pH 7.6 (NaOH) in 4 × 12.5 mL increments to elute the bound protein. The eluant was applied onto a 5 mL HiTrap Q HP anion exchange chromatography column (Cytiva Life Sciences) equilibrated with 5 mM 4-(2-hydroxyethyl)-1-piperazineethanesulfonic acid (HEPES), 50 mM citric acid, 10% glycerol, pH 7.6 (NaOH) (low-salt buffer) then washed with 50 mL of low-salt buffer before employing a 0–60% linear gradient of increasing 5 mM HEPES, 50 mM citric acid, 1 M NaCl, 10% glycerol, pH 7.6 (NaOH) (high-salt buffer) over 60 mL while fractionating into 3 mL at a flow rate of 2 mL/min. In most cases, the protein eluted at ≈0.3 M NaCl; in other cases, the protein eluted during the wash step. Protein purity was assessed by SDS-PAGE, and purified protein fractions were pooled and concentrated in 30 kDa molecular weight cutoff (MWCO) Amicon Ultra Centrifugal Filters. Protein concentrated to ≤1 mL was injected onto a 120 mL Superdex 200 pg 16/600 column (Cytiva Life Sciences) for size-exclusion chromatography (SEC) and eluted isocratically with 0.1 M citric acid, 0.1 M NaCl, 10 mM HEPES, pH 7.2 (NaOH) (SEC buffer) while collecting 3 mL fractions at a flow rate of 1 mL/min. For protein samples that were subjected to single-particle cryo-EM analysis, the same buffer sans 0.1 M NaCl was used during SEC purification, and fractions were collected manually to avoid undesired aggregated forms of the protein that eluted around 45–50 mL (Fig. S4B). Proteins purified by SEC were concentrated to ≈5–10 mg/mL in Amicon Ultra Centrifugal Filters (30 kDa MWCO) and, if not used immediately for experimentation, flash-frozen in ≤100 µL aliquots by immersion in LN2 and stored at -80 °C.

**General DBA Crosslinking.** Prior to use in crosslinking reactions, neat DBA (2 mL) was filtered through a small plug (≈0.5 mL) of anhydrous aluminum oxide in a Pasteur pipette under N_2_ gas and diluted to 0.5 M in anhydrous DMF for storage at -80 °C. This 0.5 M DBA stock solution was diluted to 5 mM in anhydrous DMF and further diluted to 0.25 mM in 50% aqueous DMF to create the working stock solution for all crosslinking reactions. The working 0.25 mM DBA stock solution was always prepared fresh in small quantities (50–100 µL) and used within 15 minutes.

**DBA Crosslinking of Wild-type and Variants of RIFS M1-DD and M1-TEII.** *Holo-* or *apo-*form RIFS M1-DD or RIFS M1-TEII harboring wild-type or mutated/altered sequences were purified according to the above procedure, concentrated to ≈7 mg/mL, and incubated individually at 5 µM with 0.1 mM TCEP in 300 mM citric acid, 20 mM HEPES, pH 7.3 (NaOH) for 15 min before adding 15 µM DBA or an equal volume of 50% aqueous DMF (control). (Note: all concentrations reflect their final concentrations after addition of DBA to the 10 µL reaction.) Crosslinking was quenched after 30 s by addition of an equal volume of 2x Laemmli buffer supplemented with 50 mM DTT and heated at 95 °C for 2 min prior to SDS-PAGE analysis using NuPAGE 3–8% Tris-Acetate Mini Protein Gels (Invitrogen).

**Preparation of Crosslinked RIFS M1-TEII for Single-particle Cryo-EM Analysis.** To isolate crosslinked M1-TEII for downstream single-particle cryo-EM analysis, we prepared multiple identical DBA crosslinking reactions and pooled them after quenching. This approach was necessary to avoid previously noted mixing effects at increased reaction volume that reduced crosslinking efficiency(5). A total of 20 × 55 µL reactions were carried out as above with slight adaptations. Specifically, *holo*-form M1-TEII was incubated at 12 µM concentration with 0.2 mM TCEP in 300 mM citric acid, 20 mM HEPES, pH 7.3 (NaOH) for 15 min before adding 24 µM DBA (total M1-TEII = 13.2 nmol). (Note: all concentrations reflect their final concentrations after addition of DBA to the 55 µL reaction.) Reactions were quenched after 30 s by the addition of 4 mM β-mercaptoethanol and buffer exchanged into SEC buffer using 7 kDa MWCO Zeba spin desalting columns (Thermo Fisher Scientific). The crosslinked M1-TEII material was pooled and used immediately for complexation with F_ab_ 1B2 and cryo-EM analysis (see **Cryo-EM Sample Preparation and Data Collection**).

**Isolation of Crosslinked and Un-crosslinked Module + F_ab_ 1B2 Complexes for Cryo-EM Analysis.** RIFS modules (M1-DD and M1-TEII) and F_ab_ 1B2 were individually purified via SEC as above prior to re-purification of the module-F_ab_ complexes. Module-F_ab_ complexes were prepared by adding 1.5 equivalents of 1B2 heterodimer per equivalent of PKS monomer, in accordance with the binding stoichiometry(6), and incubated on ice for 30 min before SEC purification of the complex (see **Protein Expression and Purification** and Fig. S4).

**Cryo-EM Sample Preparation and Data Collection.** Crosslinked and un-crosslinked module-F_ab_ complexes were concentrated to ≈10 mg/mL using Amicon Ultra Centrifugal Filters (30 kDa MWCO) before adding 0.03% nonyl phenoxypolyethoxylethanol (NP-40), 0.8 mM NADPH, and 0.8 mM TCEP, and applying 3 µL onto glow-discharged 300-mesh R 2/1 Quantifoil copper grids. The grids were blotted for 4 s at 4 °C and 100% relative humidity and vitrified in liquid ethane using a Vitrobot Mark IV (Thermo Fisher Scientific). The vitrified samples were imaged at 300 or 200 kV accelerating voltage with a Krios G3i or Glacios transmission cryo-electron microscope, respectively (Thermo Fisher Scientific). The Krios G3i was equipped with a K3 direct-electron detector (DED) and BioQuantum energy filter (Gatan), whereas the Glacios was equipped with a Falcon4 DED (Thermo Fisher Scientific) and no energy filter. The data were collected at nominal magnifications of 81,000× (Krios G3i) or 130,000× (Glacios), corresponding to a calibrated sampling of 1.1 Å/pixel or 0.73 Å/pixel, respectively. EPU software (Thermo Fisher Scientific) was used to record dose-fractionated movies in non-gain normalized .tiff format with a total dose of 50 electrons and dose rates of 10.58 or 6.65 e^–^·pixel^-1^·s^-1^ on the Krios G3i or Glacios, respectively (Table S2).

**Single-particle Cryo-EM Image Processing and 3D Reconstruction.** For details associated with cryo-EM data processing, see Figs. S5 and S10. For all datasets, dose-fractionated movies were applied to motion correction, dose weighting, and contrast transfer function (CTF) estimation in cryoSPARC(7). Additional jobs performed in cryoSPARC included particle picking, particle extraction, 2D classification, *ab initio* reconstruction, and homogenous refinement. Relion(8) was used for 3D classification by converting cryoSPARC particles in .cs format into .star format via the csparc2star.py script in pyem(9). In each case, *ab initio* models generated from ≈10–20% of the total curated micrographs were used to create templates and perform reference-based particle picking from the entire sets of curated micrographs. The particles of select classes were imported to EMAN2 and subjected to Gaussian mixture model-based orientation refinement, as well as patch-by-patch refinement, improving the resolution of flexible domains(10) (Figs. S7 and S12). C1 symmetry was specified for all 3D reconstructions and refinements.

**Model Building and Refinement.** Atomic models for individual domains (DH°, KR, and CP) and didomains (KS-AT) of RIFS M1-DD and M1-TEII and their associated docking or thioesterase domains, respectively, were generated in AlphaFold 3 and fit as rigid bodies into their corresponding cryo-EM maps using ChimeraX(11). Linker regions with supporting map density were built manually in Coot(12), and the entire map/model combinations were automatically refined in Phenix(13) using *Real-space Refinement*(14)*.* The 4′-phosphopantetheine (Ppant) cofactor was modeled by substituting the Ser residue that becomes 4′-phosphopantetheinylated with a 4HH residue in Coot. The DBA crosslink could not be observed in any of the cryo-EM maps associated with crosslinked M1-TEII and was therefore not modeled.

**Single-Turnover Kinetic Analysis of LM-M1 Catalyzed Diketide Formation by LC-MS/MS.**

Assays without pre-incubation: A 160 μL enzymatic reaction containing 5 μM *holo*-LM-M1-DD, *holo*-LM-M1-TEII*, or *holo*-LM-M1-DD(C802A) was combined with 50 mM sodium phosphate (pH 7.3, NaOH; supplemented with 1/18 equiv. HEPES), 100 mM sodium citrate (pH 7.3, NaOH; supplemented with 1/18 equiv. HEPES), 10 mM TCEP, 15 mM MgCl_2_, 10% glycerol, 1 mM sodium benzoate, 5 mM ATP, 1 mM NADPH, and 1 mM (2*R,S*)-methylmalonyl-CoA (MeMal-CoA) and incubated at 20 °C for various incubation times. After 1 min, 5 min, 15 min, 60 min, and 120 min, a 20 μL reaction aliquot was subjected to alkaline hydrolysis by adding 5 M KOH to a final concentration of 350 mM and incubating at 65 °C for 20 min (Fig. 5). Upon completion of hydrolysis, formic acid was added to a final concentration of 2.5% (ν/ν), and the hydrolysates were clarified by centrifugation at 12,000 × g for 10 min. A 10 µL aliquot of each sample was injected onto a Poroshell 120 EC-C18 column (4.6 × 100 mm, 2.7 μm, Agilent Technologies) using a 1290 Infinity II HPLC system (Agilent Technologies) and eluted at a flow rate of 600 µL/min with solvent A (aqueous 20 mM ammonium acetate) and solvent B (acetonitrile) using a 9.5-minute gradient (min/% B: 0/10, 3/100, 3.5/100, 4/10, 9.5/10). The column effluent was directed to an OptiFlow Turbo V Electrospray ionization source connected to a Triple Quad^TM^ 5500+ QTRAP 5500+ triple quadrupole mass spectrometer (AB Sciex LLC) acquiring mass spectra in targeted multiple reaction monitoring (MRM) mode in the negative polarity. In this assay, the hydrolyzed diketide (**d2′**) product precursor ion (*m/z* 179) was fragmented at a specific collision energy (-17 eV) to produce characteristic product ions (*m/z* 73 and *m/z* 117) at a specific LC retention time (2.6 min) to ensure specificity and accurate quantification in the complex biological samples (Fig. S14). Chromatographic peaks were extracted and integrated using Analyst (AB Sciex LLC). Technical replicates of each enzymatic reaction were performed in triplicate.

Assays with pre-incubation: A 140 μL enzymatic reaction containing 5 μM *holo*-LM-M1-DD or *holo*-LM-M1-TEII* was combined with 50 mM sodium phosphate (pH 7.3, NaOH; supplemented with 1/18 equiv. HEPES), 100 mM sodium citrate (pH 7.3, NaOH; supplemented with 1/18 equiv. HEPES), 10 mM TCEP, 15 mM MgCl_2_, 10% glycerol, 1 mM sodium benzoate, and 5 mM ATP and incubated at 20 °C for 150 min to allow LM-catalyzed benzoylation to reach a maximum extent (pre-incubation). Afterwards, the reaction was supplemented with 1 mM NADPH and 1 mM (2*R,S*)-MeMal-CoA to initiate diketide (**d2**) formation (Fig. 5) and further incubated at 20 °C. After 0.25 min, 0.5 min, 1 min, 5 min, 15 min, and 30 min, a 20 μL reaction aliquot was treated with KOH, heating, and formic acid in the same manner as above. The hydrolysates were clarified by centrifugation at 12,000 × g for 10 min, and the supernatants were subjected to LC-MS/MS analysis as above. Technical replicates of each enzymatic reaction were performed in triplicate.

**Quantification of LM-M1 Bound Diketide by LC-MS/MS.** Authentic standards of **d2′** were prepared at 45 nM, 450 nM, and 4,500 nM in the presence of 3.5 µM *holo*-LM-M1-DD or *holo*-LM-M1-TEII* in 50 mM sodium phosphate (pH 7.3, NaOH; supplemented with 1/18 equiv. HEPES), 100 mM sodium citrate (pH 7.3, NaOH; supplemented with 1/18 equiv. HEPES), 10 mM TCEP, 15 mM MgCl_2_, and 10% glycerol (reaction buffer). Enzymatic reactions (30 µL) were prepared separately by incubating 3.5 µM *holo*-LM-M1-DD or *holo*-LM-M1-TEII* in the above reaction buffer for 15 min at 20 °C prior to addition of 1 mM sodium benzoate, 5 mM ATP, 1 mM NADPH, and 1 mM (2*R,S*)-MeMal-CoA. Reactions were incubated for 3 h at 20 °C and terminated via buffer exchanging into reaction buffer using 7 kDa MWCO Zeba spin desalting columns (Thermo Fisher Scientific) to remove unreacted substrates. (This protocol was originally designed to enable simultaneous detection of protein-bound benzoyl and diketide (**d2**) thioester groups. However, due to complications associated with benzoyl group quantification, the buffer exchange step was later deemed unnecessary for diketide quantification.) The protein recovery following buffer exchange was determined by the Bradford assay and protein absorbance based on the calculated molecular weights and extinction coefficients. Each sample, including the above standards and enzymatic reactions, was subjected to alkaline hydrolysis by adding 5 M KOH to a final concentration of 350 mM and incubating at 65 °C for 20 min (Fig. 5). Upon completion of hydrolysis, formic acid was added to a final concentration of 2.5% (ν/ν), and the hydrolysates were clarified by centrifugation at 12,000 × g for 10 min. The supernatants were then subjected to LC-MS/MS analysis as above with a slightly modified 9.5-minute gradient (min/% B: 0/10, 5/100, 6/100, 6.1/10, 9.5/10). Data corresponding to the **d2′** standards were used to generate linear standard curves (Fig. S14) from which **d2′** concentrations were calculated in the above enzymatic reactions. Note: the reported concentrations of **d2′** in Fig. S14B reflect their final concentrations after buffer exchange and sample workup. Technical replicates of each **d2′** standard and enzymatic reaction were performed in triplicate.

**Supporting Figures**

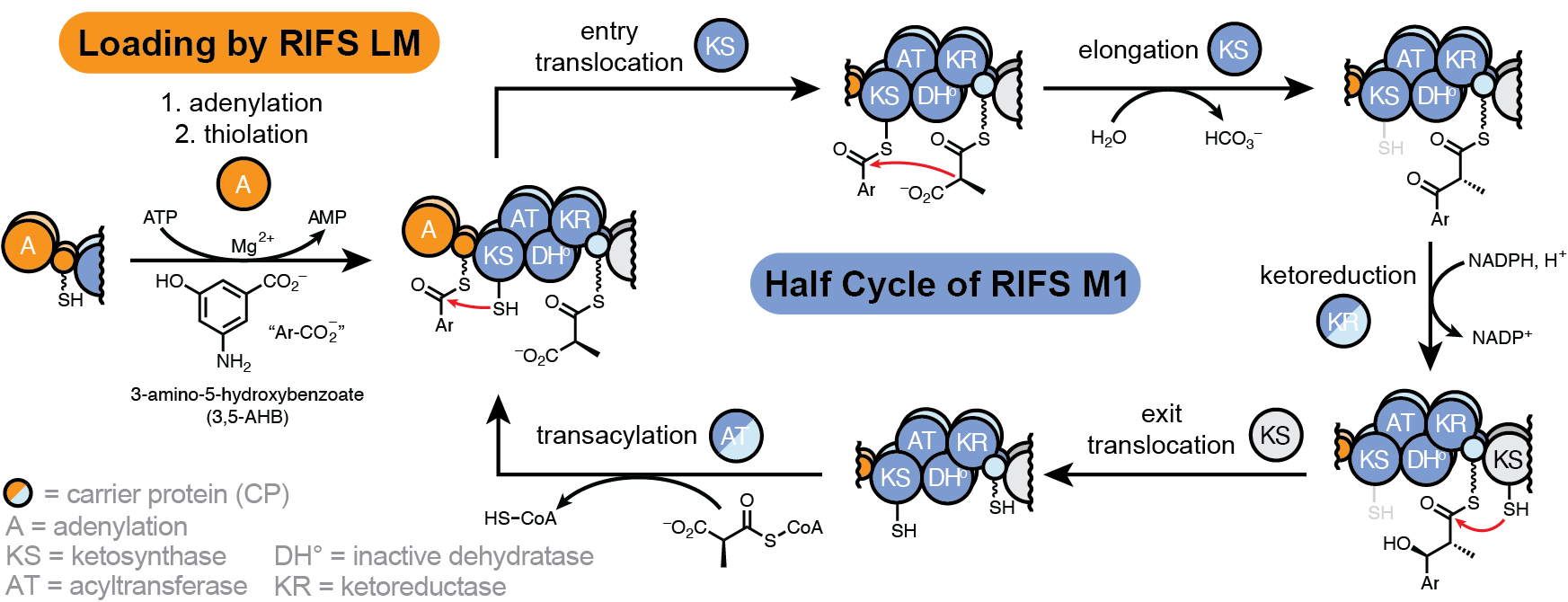

**Fig. S1.** Catalytic cycle of one half of the rifamycin synthetase (RIFS) loading module (LM) + module 1 (M1). The adenylation (A) domain catalyzes two-step 3-amino-5-hydroxybenzoylation of its loading carrier protein (CPL) domain, in orange. The first step involves ATP-dependent adenylation of 3-amino-5-hydroxybenzoate (3,5-AHB) into 3-amino-5-hydroxybenzoyl-AMP. The second step involves transfer of the activated 3-amino-5-hydroxybenzoyl group onto the 4′-phosphopantetheine (Ppant) cofactor of CPL. The ketosynthase (KS) domain of the downstream PKS module (M1) then catalyzes transfer of the 3-amino-5-hydroxybenzoyl group onto its own catalytic Cys residue (entry translocation). (It was previously observed that this transfer only occurs after acyltransferase (AT)-catalyzed transacylation of 2*S*-methylmalonyl (MeMal) from MeMal-CoA onto the Ppant cofactor of M1’s CP domain (CP1), in light blue(15).) Attachment of both acyl groups onto M1 primes KS-catalyzed decarboxylative Claisen condensation between the MeMal nucleophile and KS-bound electrophilic thioester, resulting in a β-ketoacyl-CP1 thioester product (elongation). The ketoreductase (KR) domain catalyzes stereospecific ketoreduction to the (2*R*,3*S*) diastereomer of the diketide, which is then received by the downstream KS from RIFS module 2 (M2) (exit translocation). The dehydratase domain of M1 is inactive (denoted DH°), presumably due to presence of a Gly in place of the conserved catalytic His (position 1521 in RifA, WP_013222547.1)(16). This illustration depicts the activities of a single catalytic subunit defined as the complete set of enzymes required for one catalytic half-cycle. However, it does not necessarily reflect accurately which of the two catalytic subunits are employed from one half-cycle to the next, nor does it reflect the possibility that activities on both subunits happen concurrently. Note: in accordance with previous evidence, heavy and light shading of the two subunits was used to illustrate *intra*-molecular entry/exit translocation and *inter-*molecular elongation (3, 17). The KS Cys is shown in black and gray to indicate active an inactive states, respectively, based on the ‘turnstile’ mechanism(18). That is, after KS-catalyzed elongation, the KS Cys becomes transiently inactivated until exit translocation of the polyketide intermediate.

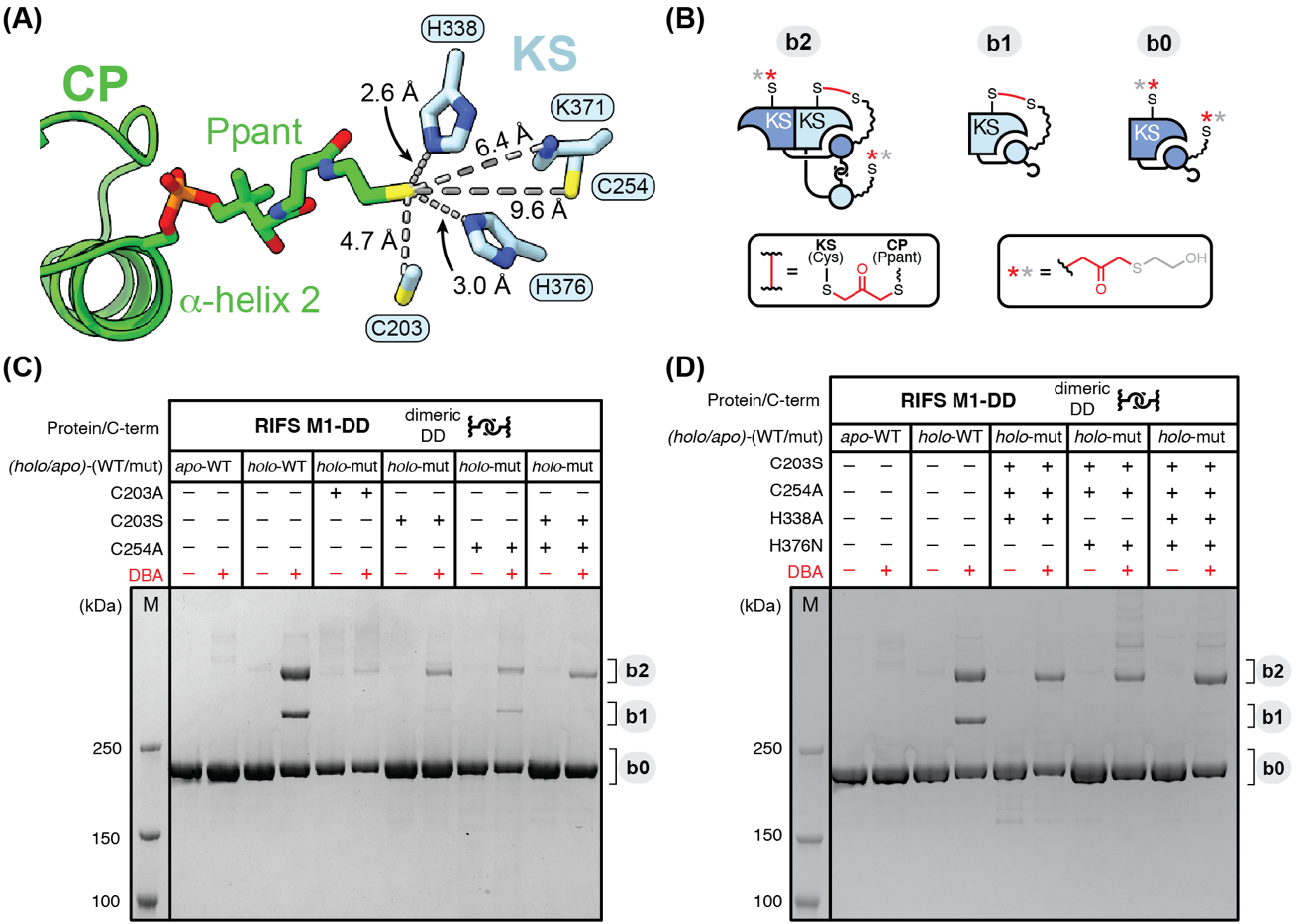

**Fig. S2.** Site-selectivity of DBA crosslinking was investigated through mutational analysis of RIFS M1. **(A)** The structure of RIFS M1 in the *elongation mode* (PDB 9PAV) was used to guide the selection of nucleophilic residues for mutation and analysis of their contributions to DBA crosslinking. **(B)** Cartoon depictions of the three major products following addition of DBA to RIFS M1. **(C)** Wild-type (WT) or mutated (mut) RIFS M1 in *holo*- or *apo*-forms were subjected to DBA crosslinking (or DMF as a control) in the presence of 0.1 mM TCEP, 20 mM HEPES, and 300 mM citrate (pH 7.3), followed by reducing SDS-PAGE analysis. Only trace amounts of non-specific products, relative to starting materials, were observed in the *apo*-WT M1 reactions, indicating that the Ppant thiol group of the carrier protein domain was essential for formation of the major crosslinked products observed in *holo*-WT M1: i.e., bands 1–2 (**b1**–**b2**). Mutation of the catalytic Cys residue of the KS domain to Ala (C203A) or Ser (C203S) reduced crosslinking by approximately 8-fold and 5-fold, respectively, suggesting that **b1** and **b2** are principally comprised of Ppant-Cys203 crosslinks. A similar reduction was observed when Cys254, which is positioned ~10 Å away from the Ppant in the cryo-EM structure, was mutated to Ala (C254A). Combining two of these mutations (C203S/C254A) indicated that other nucleophilic residues besides Cys203 and Cys254 are likely to partake in crosslinking with Ppant. The near complete loss of **b1** in the crosslinked C203A, C203S, and C203S/C254A proteins implied that this product is mainly dependent on Cys203. **(D)** We considered that residual formation of **b2** after removal of Cys203 and/or Cys254 was due to crosslinking between Ppant and active site His residues (His338 and His376) proximal to Cys203. Hence, we created C203S/C254A/H338A and C203S/C254A/H376N triple mutants and a C203S/C254A/H338A/H376N quadruple mutant for similar DBA crosslinking. Surprisingly, although **b1** is nearly abolished, the crosslinking yields of **b2** achieved by these mutants relative to the mutants in panel C were not diminished, but rather enhanced. Together, we speculate that these triple and quadruple mutants were impaired in their ability to dock Ppant in the KS active site, thus promoting other as-yet unidentified Ppant-nucleophile crosslinks (e.g., Ppant-Ppant crosslink between both subunits). While these experiments established that other nucleophiles besides Cys203 can crosslink with Ppant, only the Ppant-Cys203 major crosslink is depicted in the graphical illustration for simplicity.

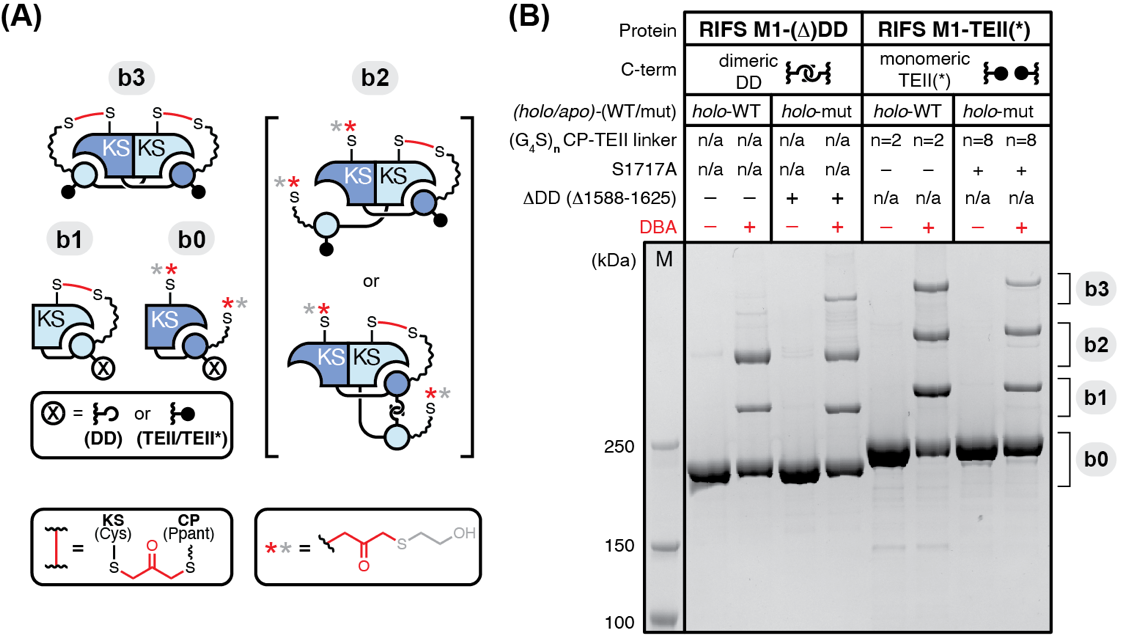

**Fig. S3.** DBA crosslinking analysis of RIFS M1-DD, M1-ΔDD, M1-TEII, and M1-TEII* (the asterisk implies S1717A and a (G_4_S)_8_ M1-TEII linker). **(A)** Cartoon depictions of the four major products following addition of DBA to these proteins. **(B)** DBA or DMF (control) was added to the above proteins in the presence of 0.1 mM TCEP, 20 mM HEPES, and 300 mM citrate (pH 7.3) and analyzed by reducing SDS-PAGE. The appearance of a band distribution in M1-ΔDD like that of M1-TEII pointed to an inhibitory role for DD in the formation of **b3**. Similar crosslinked products (**b1**–**b3**) observed in M1-TEII and M1-TEII* indicated that the linker between the CP and TEII domains (i.e., S(G_4_S)_2_ vs. S(G_4_S)_8_, respectively) was not a major determinant of crosslinking, nor was the active site Ser of the TEII (i.e., S1717 which is mutated to Ala in M1-TEII*).

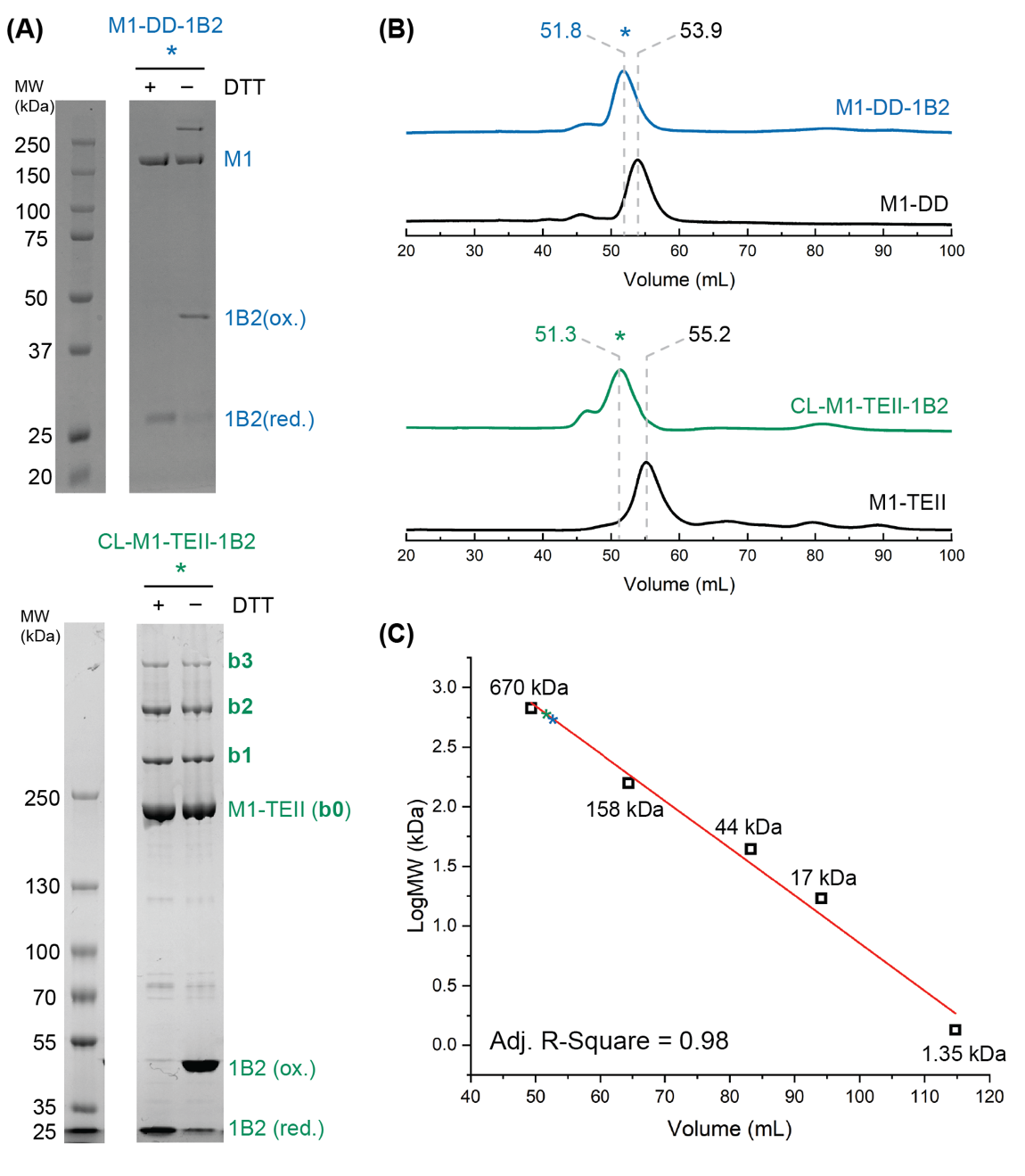

**Fig. S4.** SDS-PAGE and size-exclusion chromatography (SEC) analysis of each cryo-EM sample used in this study. **(A)** SDS-PAGE analyses of the M1-DD-1B2 and CL-M1-TEII-1B2 complexes under reducing and non-reducing conditions (± 25 mM DTT). **(B)** SEC chromatograms of RIFS M1-DD or M1-TEII alone (black) and in complex with F_ab_ 1B2 (blue and green, respectively). Note: unlike M1-DD, M1-TEII was crosslinked prior to addition of 1B2 (see **Preparation of Crosslinked RIFS M1-TEII for Single-particle Cryo-EM Analysis)**. **(C)** A semi-log standard curve generated by similar SEC analysis of protein standards (Bio-Rad #1511901).

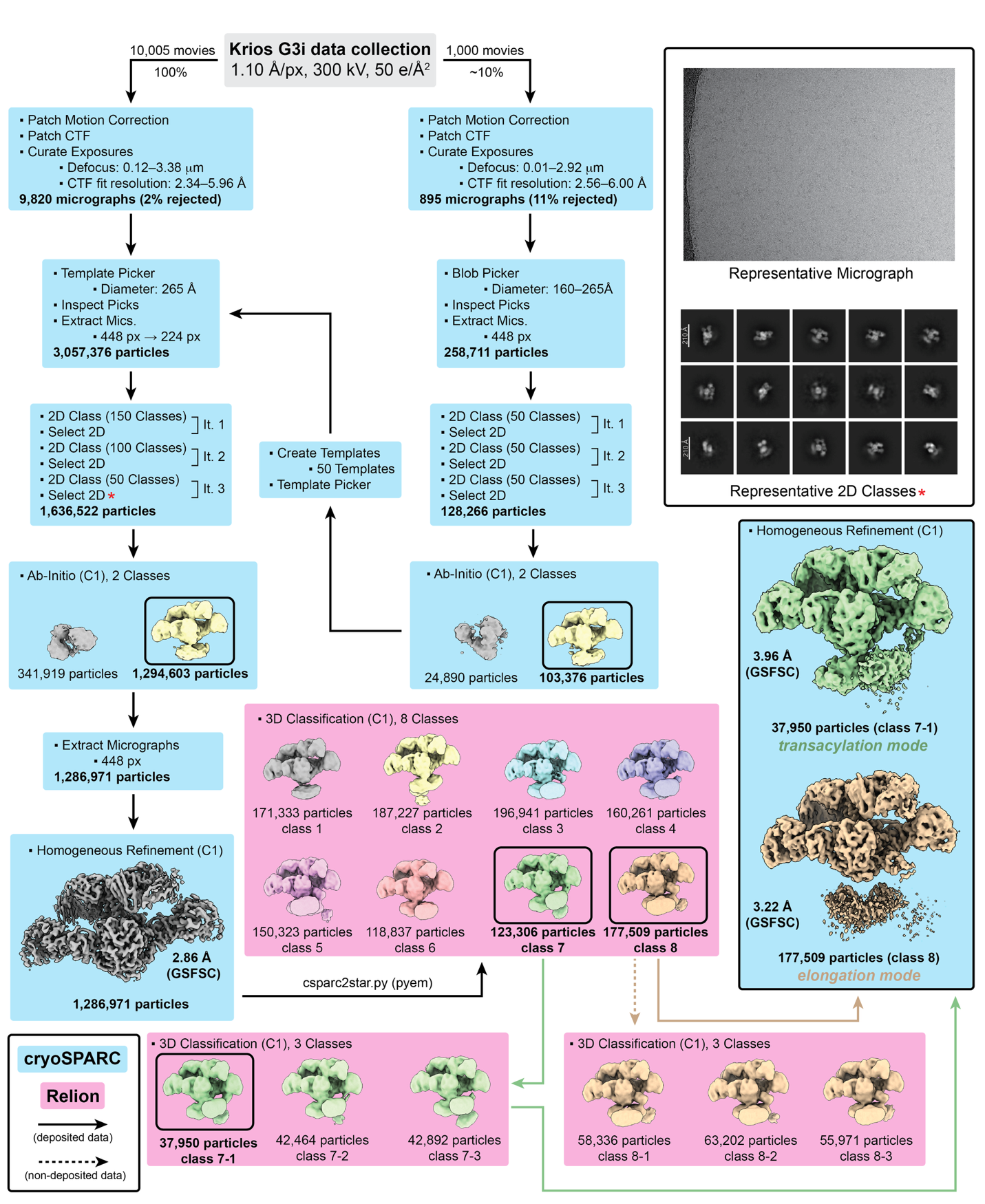
**Fig. S5.** Workflow of single-particle cryo-EM analysis of M1-DD-1B2 (as prepared in Fig. S4). A combination of processing tools in Relion(8) (pink) and cryoSPARC(7) (blue) were implemented to obtain the final cryo-EM maps (boxed).

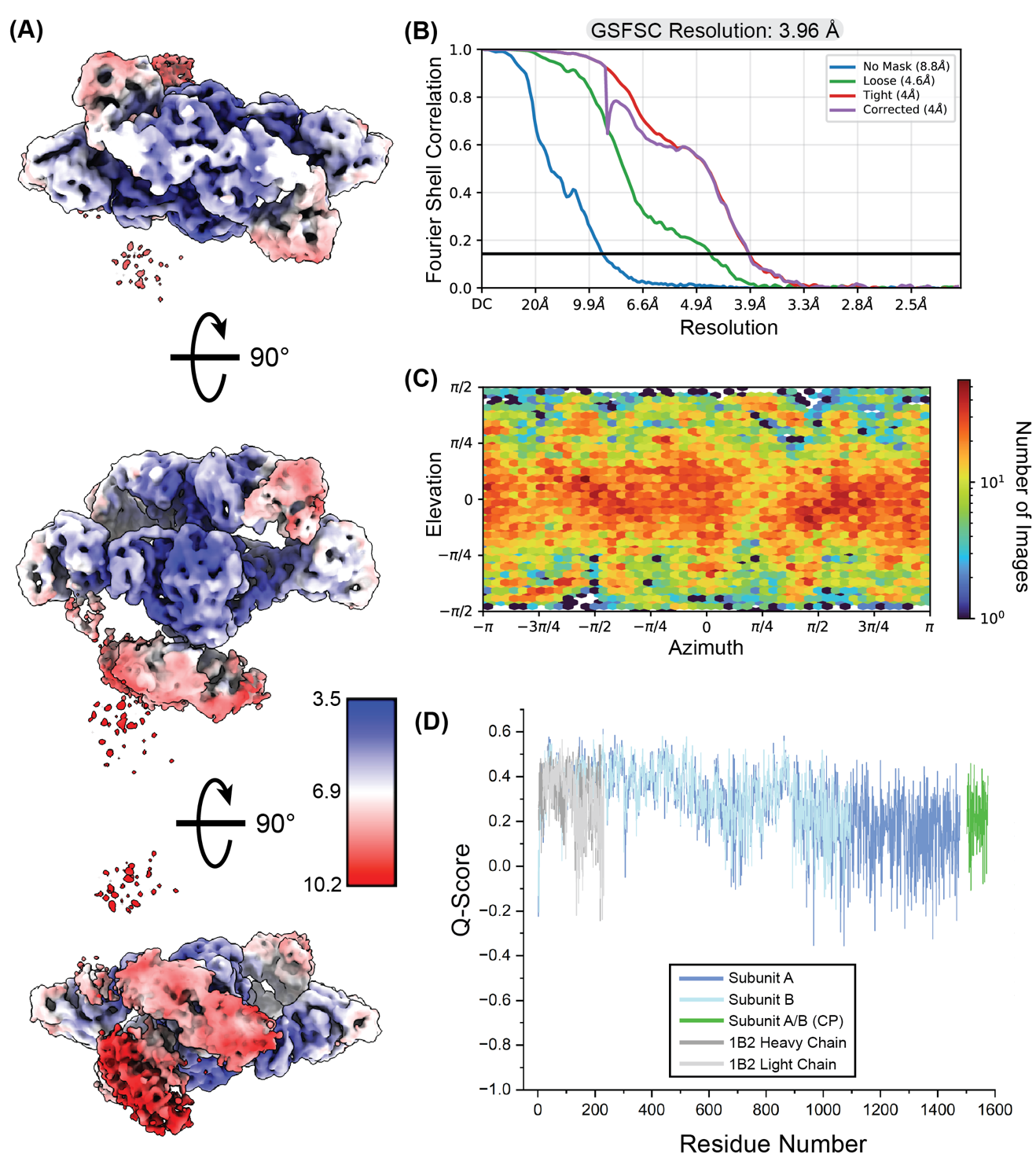

**Fig. S6.** Cryo-EM map and model validation of the *transacylation-mode* structure of M1-DD-1B2 (Fig. S5). (**A**) Local resolution map generated in cryoSPARC(7). (**B**) Curves display the Fourier shell correlation (FSC) between two independently refined half maps at different spatial frequencies generated in cryoSPARC(7). The gold-standard FSC (GSFSC) resolution is defined as the spatial frequency at which the corrected FSC curve intersects with a threshold FSC value of 0.143(19). (**C**) Euler angle distribution plot generated in cryoSPARC shows a heat map representation of the number of images for each particle orientation(7). (**D**) The Q-score assessment of atom resolvability within the cryo-EM map is plotted for all residues of the *transacylation-mode* model (PDB 9PAT)(20).

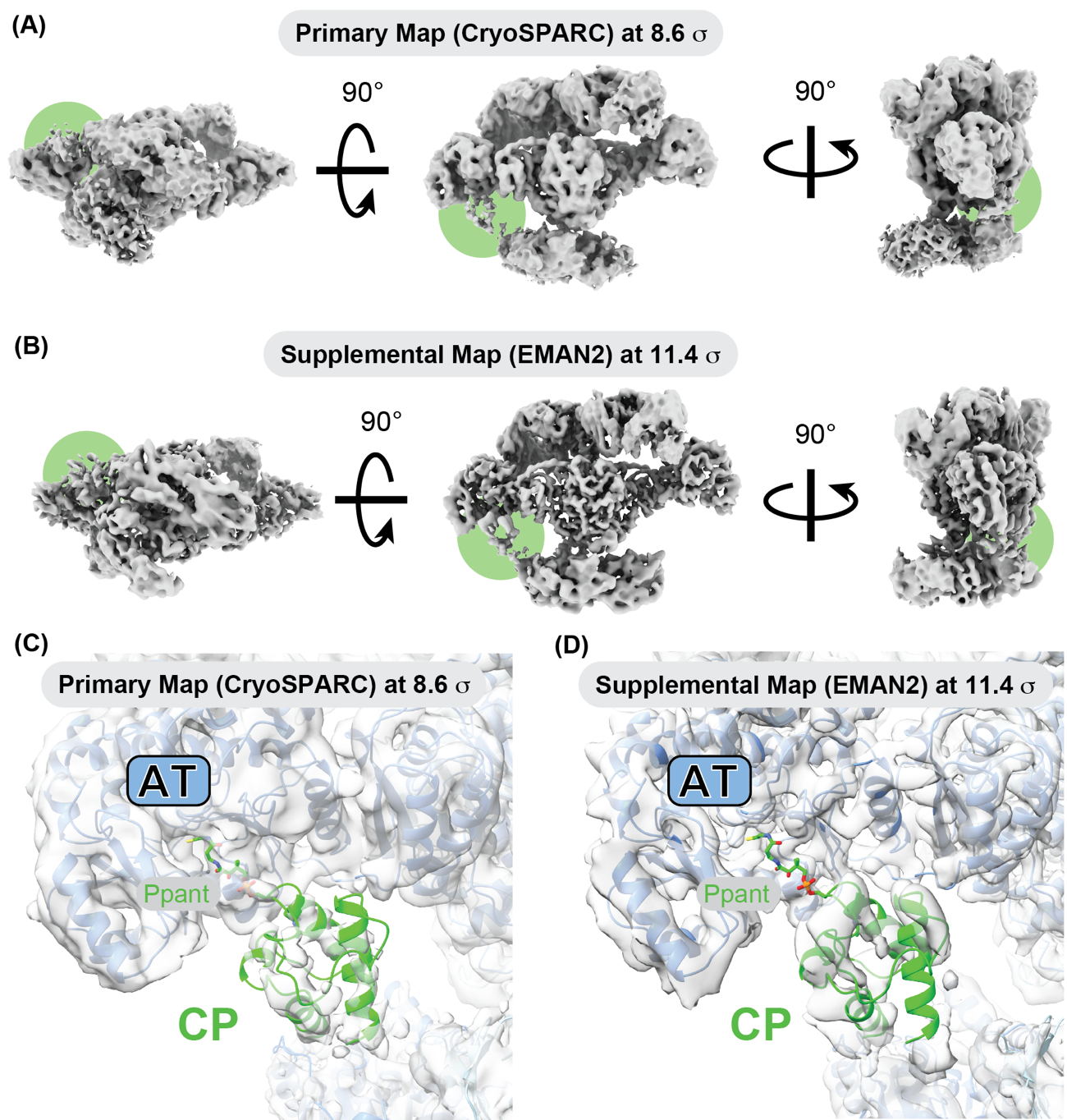

**Fig. S7.** Three different orientations of the **(A)** primary and **(B)** supplemental cryo-EM maps associated with the *transacylation-mode* structure (Figs. S5–S6) contoured at 8.6 σ and 11.4 σ, respectively. Green circles highlight regions of CP domain density. **(C–D)** Close-up views of the CP-bound AT active site cleft for the **(C)** primary and **(D)** supplemental maps highlights stronger signal for the CP domain in the supplemental map, albeit with poor atom resolvability (Fig. S6D). Sigma values were calculated from the entire cryo-EM map volumes by the equation: σ = (threshold – mean) / standard deviation.

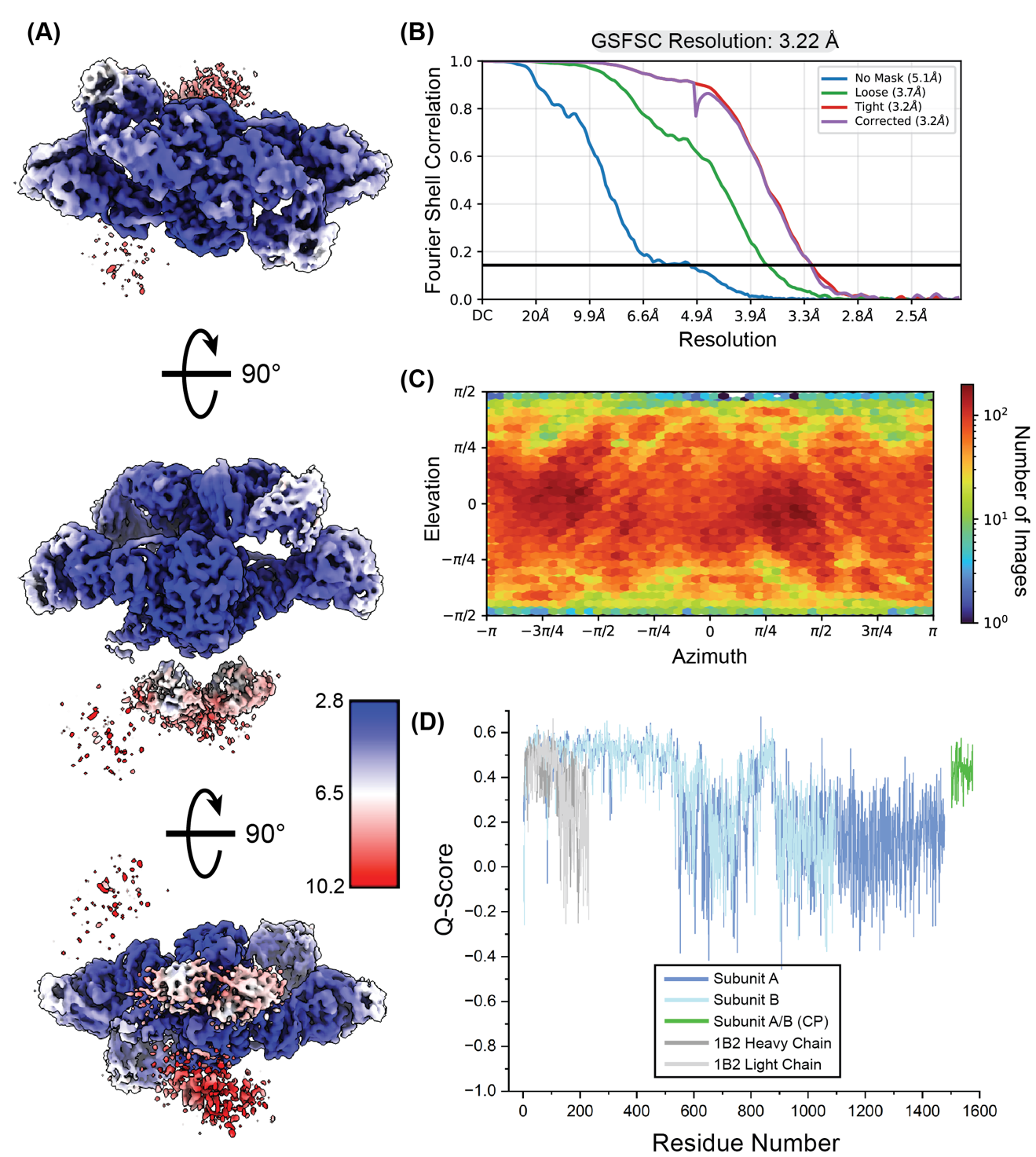

**Fig. S8.** Cryo-EM map and model validation of the *elongation-mode* structure of M1-DD-1B2 (Fig. S5). (**A**) Local resolution map generated in cryoSPARC(7). (**B**) Curves display the Fourier shell correlation (FSC) between two independently refined half maps at different spatial frequencies generated in cryoSPARC(7). The gold-standard FSC (GSFSC) resolution is defined as the spatial frequency at which the corrected FSC curve intersects with a threshold FSC value of 0.143(19). (**C**) Euler angle distribution plot generated in cryoSPARC shows a heat map representation of the number of images for each particle orientation(7). (**D**) The Q-score assessment of atom resolvability within the cryo-EM map is plotted for all residues of the *elongation-mode* model (PDB 9PAV)(20).

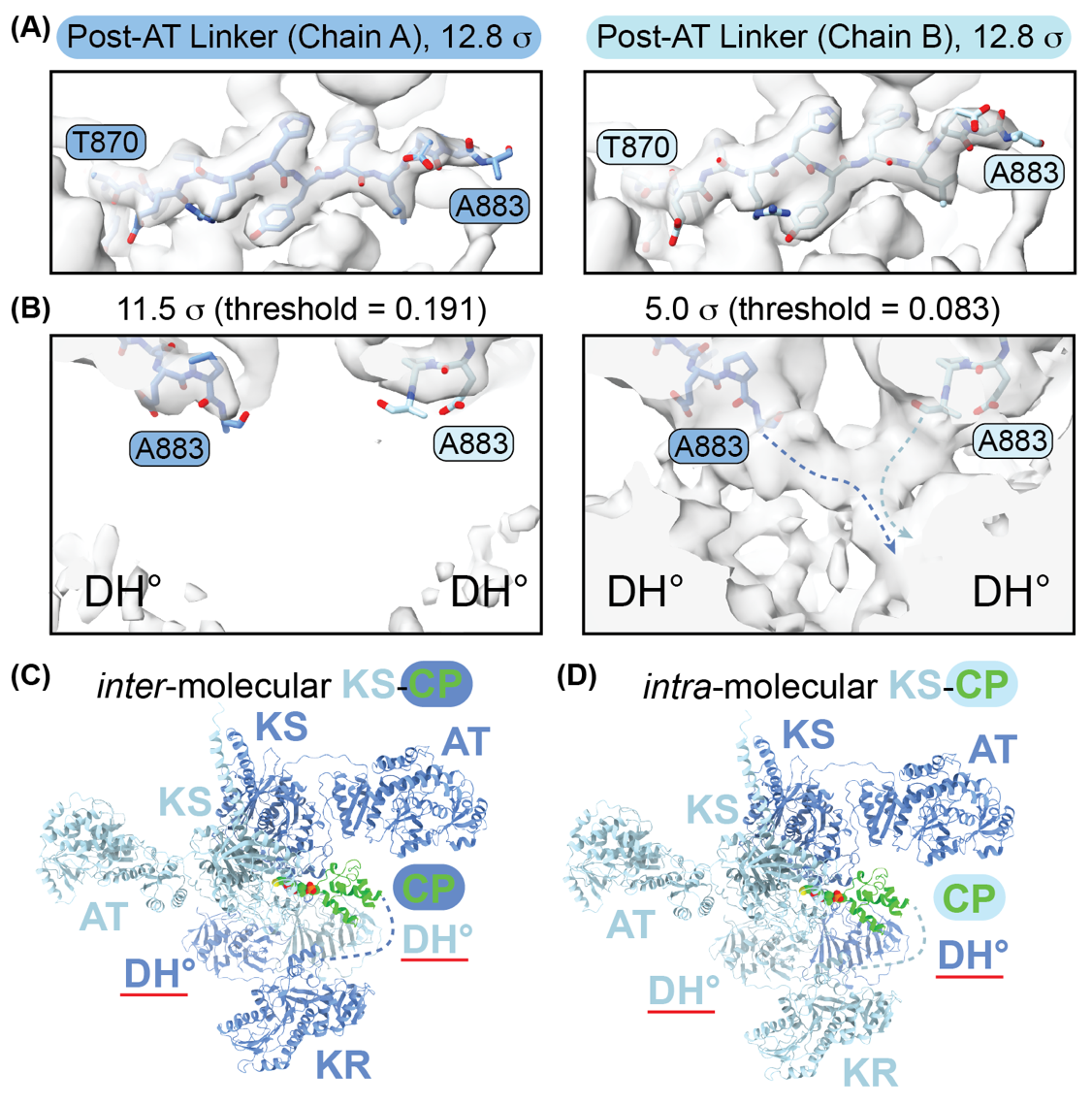

**Fig. S9.** Analysis of the inter-domain linker between the AT and DH° domains in the M1-DD-1B2 *elongation-mode* cryo-EM map (Figs. S5 and S8). **(A)** The AT and DH° domains are separated by the ‘post-AT’ linker (PAL) spanning approximately A863–G889. Residues A863–P882 of the PAL are well resolved by the cryo-EM map in both subunits. Residues T870–A883 of the PAL are shown inside the cryo-EM map contoured at 12.8 σ. **(B)** The region of density expected to harbor residues A883–G889 of the PAL in both subunits is shown at two different sigma (or threshold) values. Although the poor resolution in this region precluded accurate modeling of A883–G889, continuous density is observed at 5 σ between both PALs and a single DH° domain. This observation is consistent with two possible orientations of the DH° dimer related by a 180° rotation about the pseudo-C2 axis of module symmetry. By extension, one of them **(C)**, corresponding to the dark blue arrow in panel B, would support *inter*-molecular KS-CP interactions, whereas the other **(D)**, corresponding to the light blue arrow in panel B, would support *intra*-molecular KS-CP interactions. Although the KR-CP linker (depicted by the dashed line in panels C and D) was not well resolved by this cryo-EM map (depicted by the dashed line), we inferred based on the cryo-EM map of CL-M1-TEII-1B2 that the visible KR is connected to the visible CP (Figs. 4 and S10–S12). Sigma values were calculated from the entire cryo-EM map volumes by the equation: σ = (threshold – mean) / standard deviation.

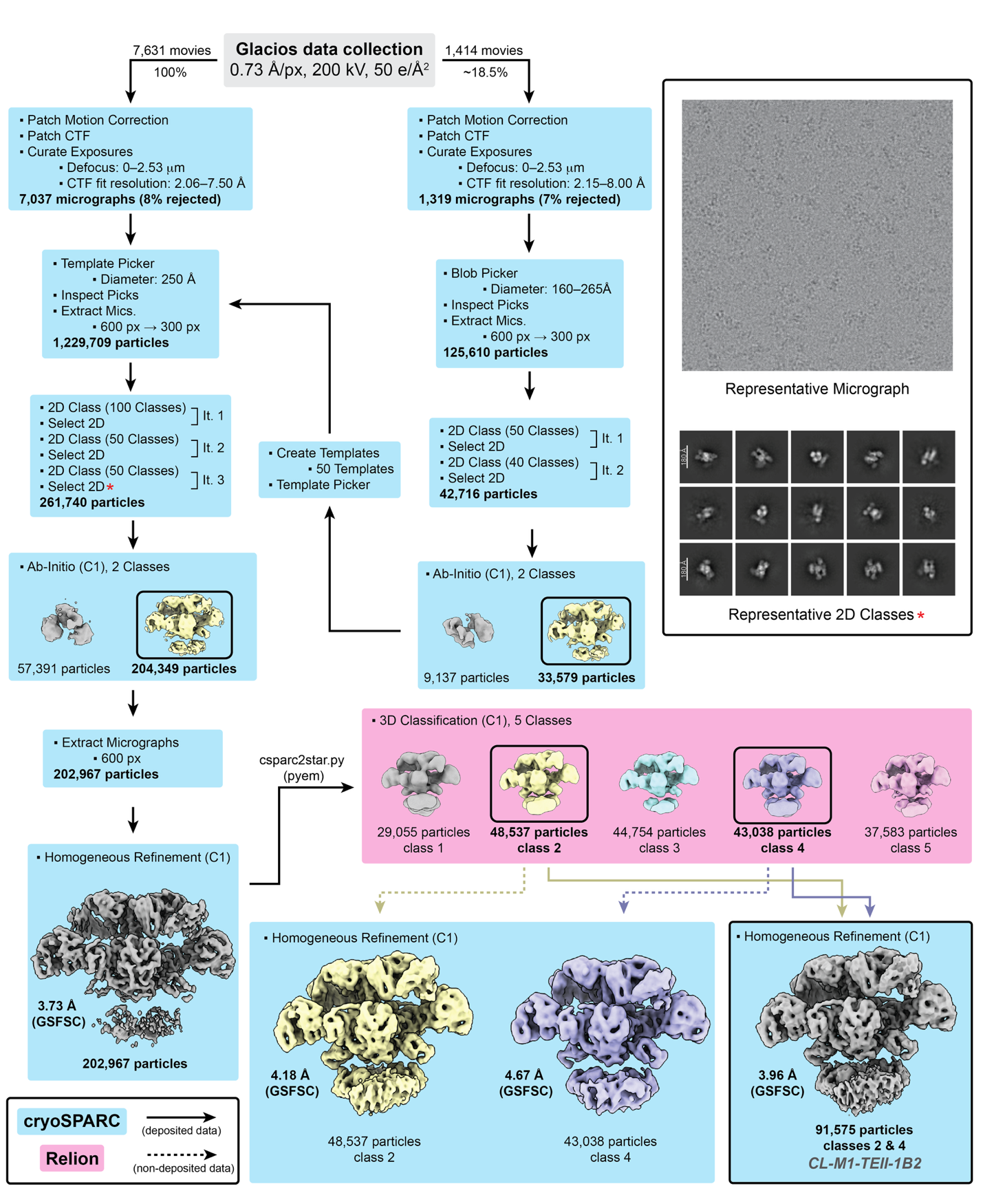

**Fig. S10.** Workflow of single-particle cryo-EM analysis of CL-M1-TEII-1B2 (as prepared in Fig. S4). A combination of processing tools in Relion(8) (pink) and cryoSPARC(7) (blue) were implemented to obtain the final cryo-EM map (boxed). Note that ‘class 3’ under 3D Classification contains density for only one CP domain, as expected for **b1** and **b2** (Figs. 2 and S4).

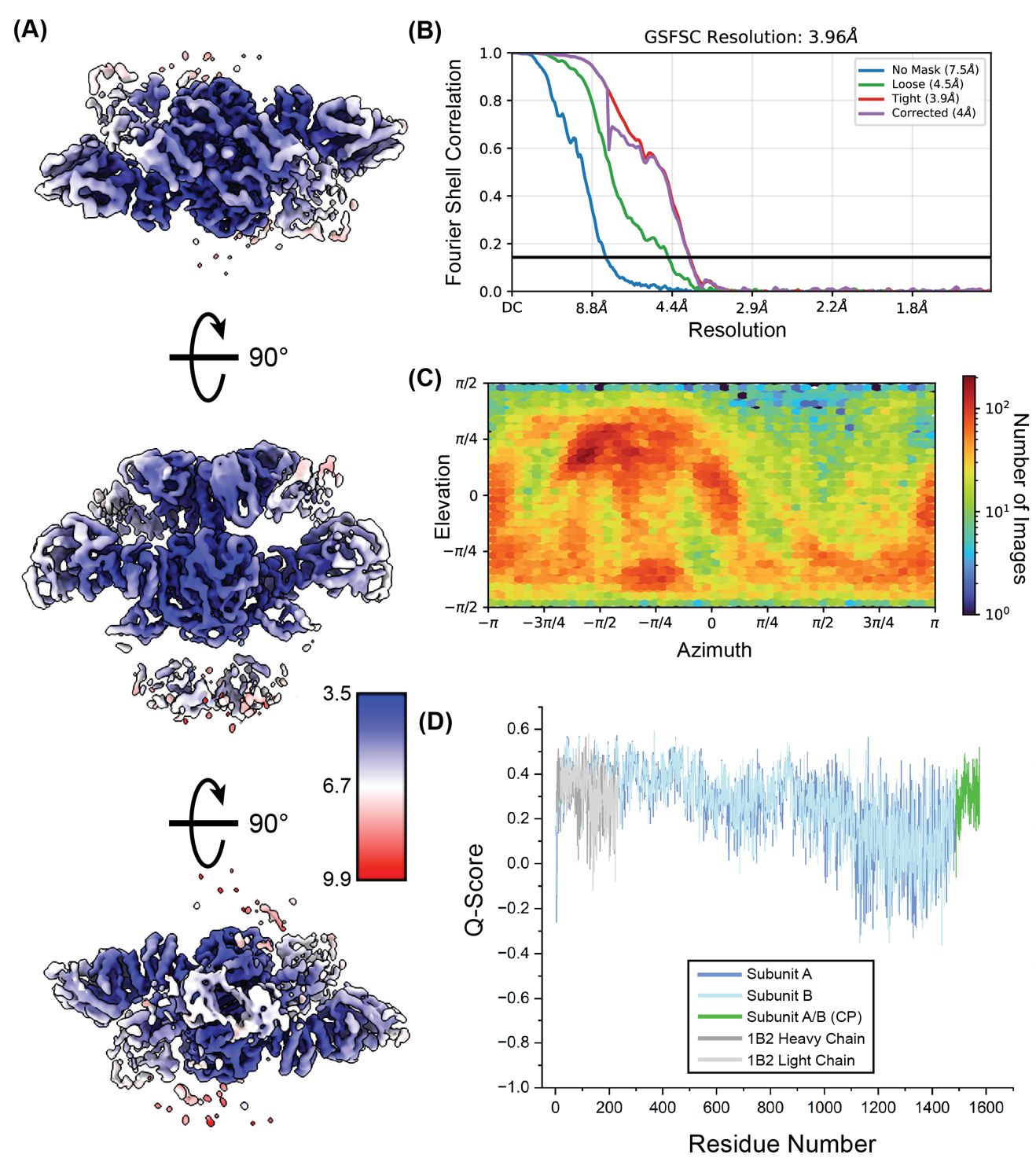

**Fig. S11.** Cryo-EM map and model validation of the structure of CL-M1-TEII-1B2 (Fig. S10). (**A**) Local resolution map generated in cryoSPARC(7). (**B**) Curves display the Fourier shell correlation (FSC) between two independently refined half maps at different spatial frequencies generated in cryoSPARC(7). The gold-standard FSC (GSFSC) resolution is defined as the spatial frequency at which the corrected FSC curve intersects with a threshold FSC value of 0.143(19). (**C**) Euler angle distribution plot generated in cryoSPARC shows a heat map representation of the number of images for each particle orientation(7). (**D**) The Q-score assessment of atom resolvability within the cryo-EM map is plotted for all residues of the CL-M1-TEII-1B2 model (PDB 9PC6)(20).

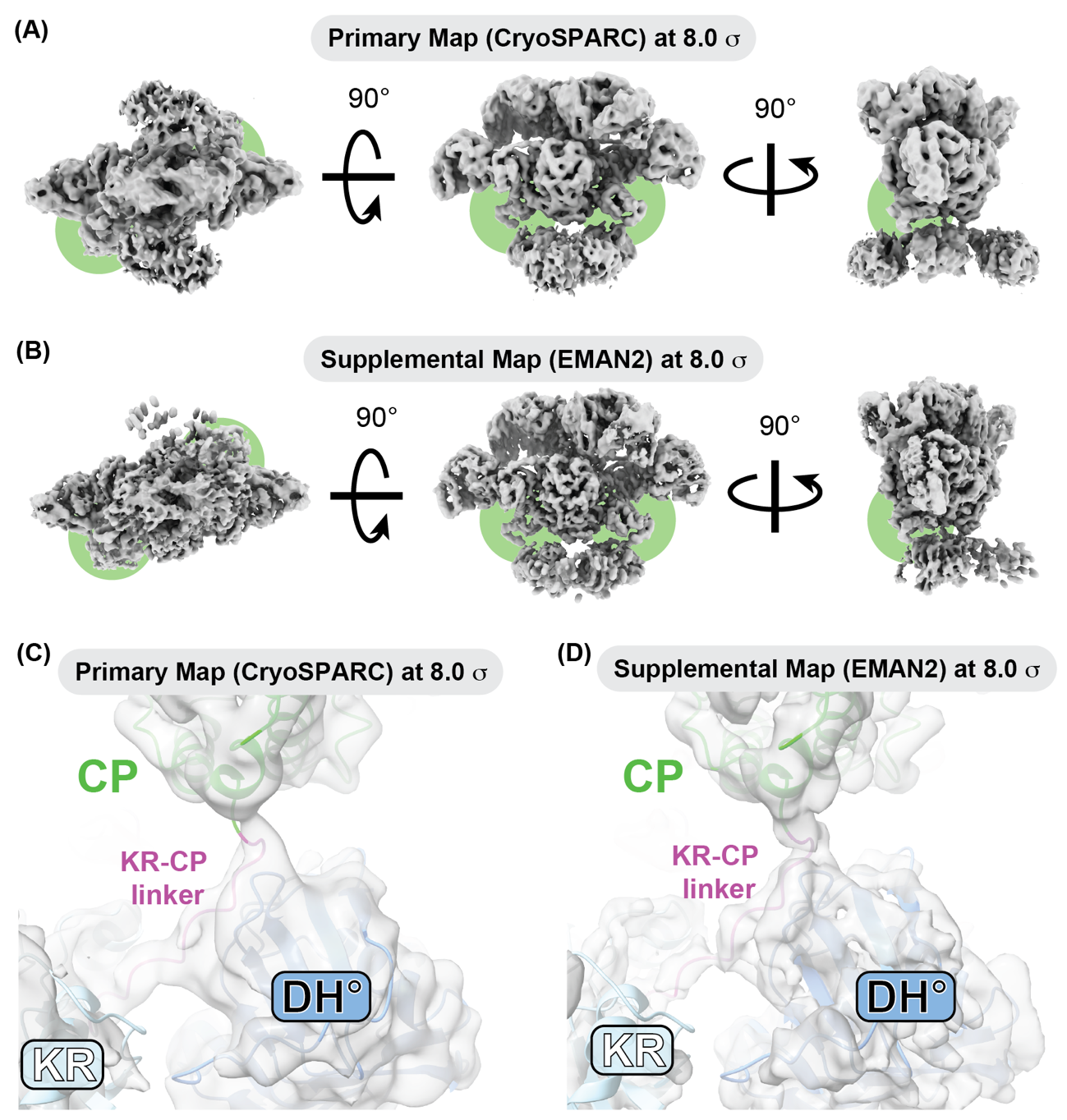

**Fig. S12.** Three different orientations of the **(A)** primary and **(B)** supplemental cryo-EM maps associated with the CL-M1-TEII-1B2 structure (Figs. S10–S11) contoured at 8.6 σ and 11.4 σ, respectively. Green circles highlight regions of CP domain density. **(C–D)** Close-up view of the KR-CP linker for the **(C)** primary and **(D)** supplemental maps highlights slightly improved local resolution in the supplemental map (Fig. S11D). Sigma values were calculated from the entire cryo-EM map volumes by the equation: σ = (threshold – mean) / standard deviation.

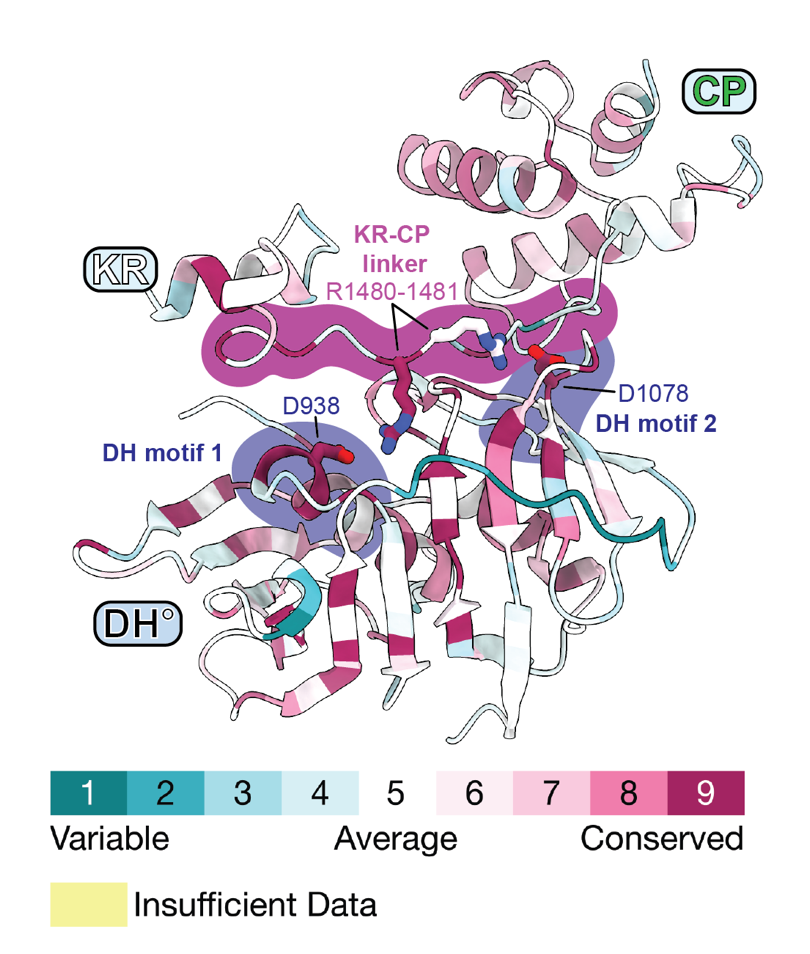

**Fig. S13.** The ConSurf server was used to represent sequence conservation as a numerical color scale on the CL-M1-TEII-1B2 model structure (PDB 9PC6)(21). Specifically, residues involved in putative inter-subunit interactions between the KR-CP linker (R1480–R1481) and DH° domain (D938 from motif 1 and D1078 from motif 2) are shown (Fig. 4B–C). Sequence conservation scores for each residue were generated from a multiple sequence alignment of 250 homologs of the DH°-KR-CP fragment of RIFS M1 (WP_013222547.1)(22).

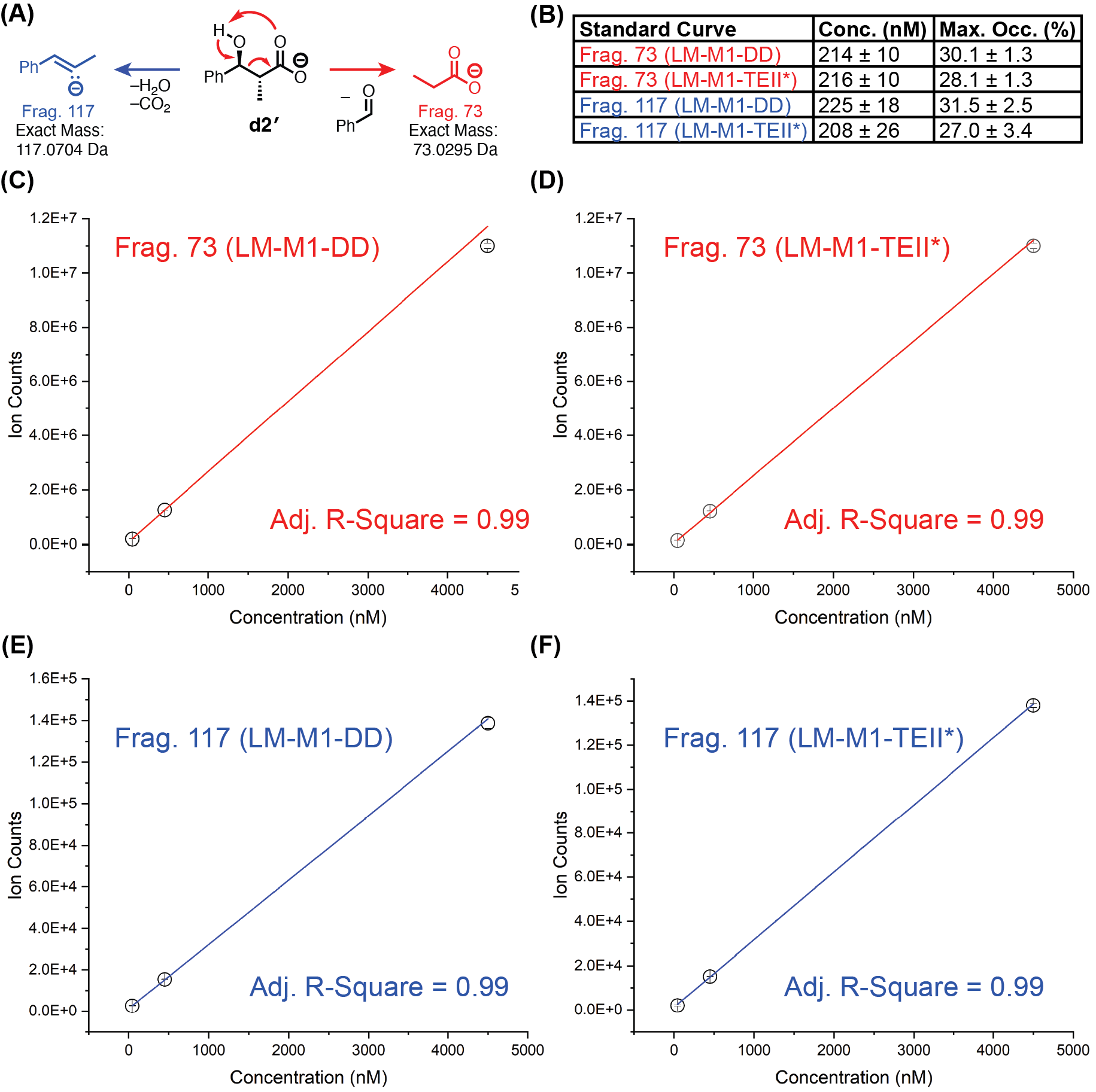

**Fig. S14.** LC-MS/MS quantitation of **d2′** using MRM. **(A)** Proposed fragmentation patterns for **d2′** based on detection of fragment ions at *m/z* 73 Da and 117 Da in negative ion mode. Standard curves for quantification of **d2′** in enzymatic reactions were generated by preparing **d2′** at 45 nM, 450 nM, and 4,500 nM in the presence of LM-M1-DD or LM-M1-TEII* in reaction buffer lacking ATP, NADPH, and MeMal-CoA. Standards were treated with KOH, heating, and formic acid in the same manner as enzymatic reactions prior to LC-MS/MS analysis. **(B)** The concentrations of **d2′** and maximum occupancy of **d2** (Fig. 5) for LM-M1-DD and LM-M1-TEII* were determined from **(C–F)** standard curves corresponding to the 73 Da and 117 Da fragment ions quantified in the presence of each protein. Note: the reported concentrations of **d2′** in panel B reflect their final concentrations after buffer exchange and sample workup (see **Quantification of LM-M1 Bound Diketide by LC-MS/MS**).

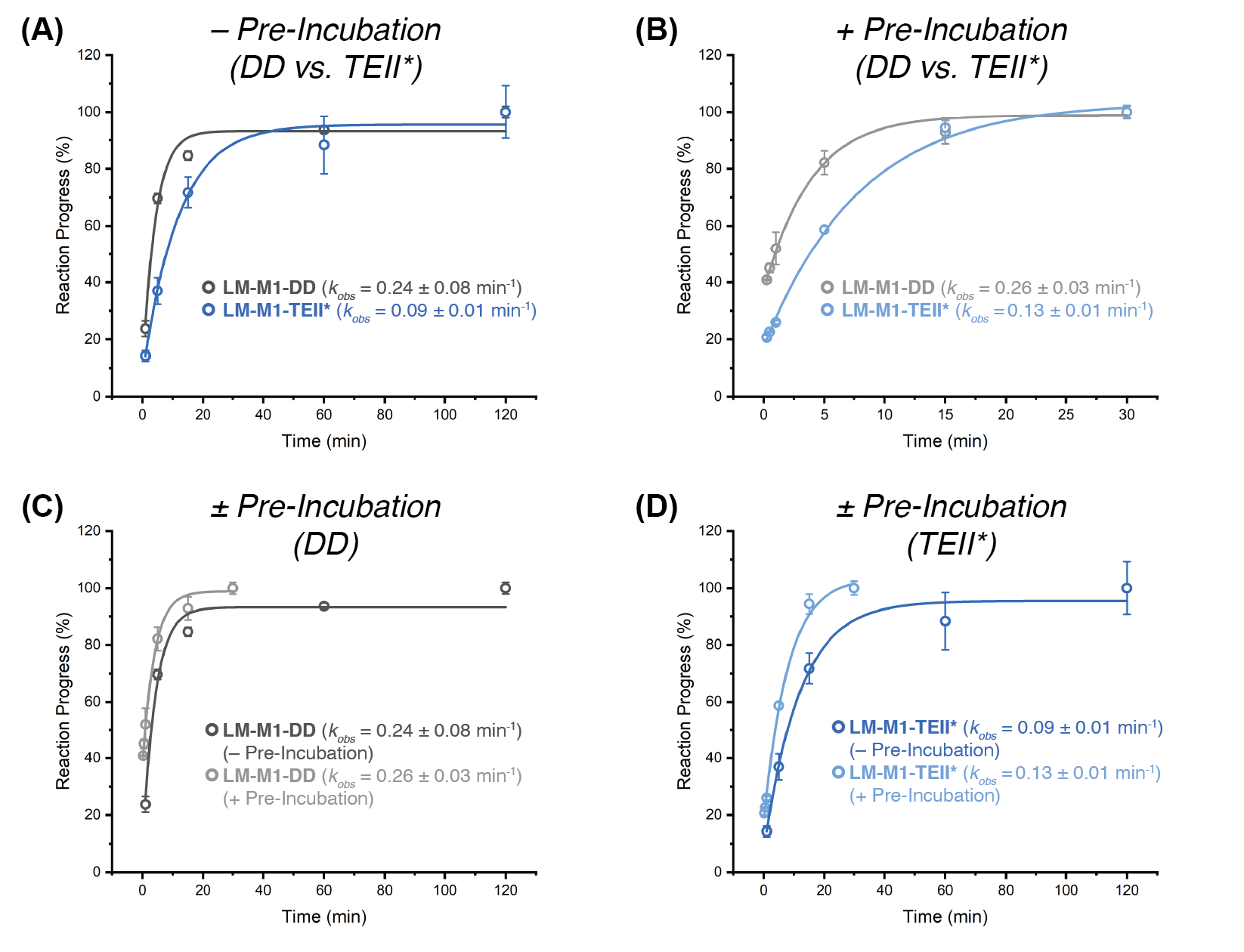

**Fig. S15.** The effect of pre-incubating ATP/Mg^2+^ and benzoate with LM-M1 harboring a C-terminal DD or TEII* on the rates of **d2** formation (Fig. 5). Black/gray traces indicate reactions with LM-M1-DD without or with pre-incubation, respectively. Blue/light blue traces indicate reactions with LM-M1-TEII* without or with pre-incubation, respectively (see **Single-Turnover Kinetic Analysis of LM-M1 Catalyzed Diketide Formation by LC-MS/MS**). Comparison of **(A)** the rates of **d2** formation by LM-M1-DD and LM-M1-TEII* without pre-incubation (replotted from Fig. 5B), **(B)** the rates of **d2** formation by LM-M1-DD and LM-M1-TEII* with pre-incubation, **(C)** the rates of **d2** formation by LM-M1-DD without or with pre-incubation, and **(D)** the rates of **d2** formation by LM-M1-TEII* without or with pre-incubation. Rate constants were obtained by fitting these data to a single-phase exponential growth function in Origin 2023b. In both cases (± pre-incubation), LM-M1-DD catalyzed **d2** formation faster than LM-DD-TEII* by a factor ≥2. Interestingly, pre-incubation had no effect on the rate of **d2** formation by LM-M1-DD, whereas it had a measurable effect on that of LM-M1-TEII*. This implied that LM-M1-TEII* was more rate-limited by LM-catalyzed benzoylation than LM-M1-DD. We note that these enzymatic reactions contained elevated amounts of citrate (100 mM), which is known to enhance the catalytic activity of PKS modules *in vitro*(3, 23, 24).

**Supporting Tables**

Table S1. Plasmids used in this study (Km = kanamycin; Cb = carbenicillin; Am = ampicillin).

| **Plasmid** | **Encoded Protein** | **Antibiotic** | **Reference** |
| --- | --- | --- | --- |
| pAM6 | RifA LM-M1-TEII(S2316A)-His_6_ (**LM-M1-TEII***) | Cb/Am | This study |
| pCL17 | RifA LM-M1-DD(C802A)-His_6_ | Cb/Am | This study |
| pDC42 | RifA LM-M1-DD-His_6_ (**LM-M1-DD**) | Cb/Am | This study |
| pDC43 | RifA M1-DD-His_6_ (**M1-DD**) | Cb/Am | This study |
| pDC85 | RifA M1-TEII-His_6_ (**M1-TEII**) | Cb/Am | This study |
| pDC147 | RifA M1-TEII(S1717A)-His_6_ | Cb/Am | This study |
| pDC155 | RifA M1-DD(Δ1588–1625)-His_6_ (**M1-ΔDD**) | Cb/Am | This study |
| pRSG56 | His_6_-Sfp | Km | (25) |
| pRW9 | RifA M1-DD(C203A)-His_6_ | Cb/Am | This study |
| pRW20 | RifA M1-DD(C254A)-His_6_ | Cb/Am | This study |
| pRW22 | RifA M1-DD(C203S/C254A)-His_6_ | Cb/Am | This study |
| pRW24 | RifA M1-DD(C203S)-His_6_ | Cb/Am | This study |
| pRW32 | RifA M1-DD(C203S/C254A/H338A)-His_6_ | Cb/Am | This study |
| pRW35 | RifA M1-DD(C203S/C254A/H376N)-His_6_ | Cb/Am | This study |
| pRW36 | RifA M1-DD(C203S/C254A/H338A/H376N)-His_6_ | Cb/Am | This study |
| n/a | F_ab_ 1B2-His_6_ (**1B2**) | Cb/Am | (6) |

**Table S2.** Cryo-EM data collection parameters.

| *Sample* | M1-DD-1B2 | CL-M1-TEII-1B2 |
| --- | --- | --- |
| *Module expression plasmid* | pDC43 | pDC85 |
| *Microscope* | Krios G3i | Glacios |
| *Voltage (kV)* | 300 | 200 |
| *Camera* | K3 | Falcon4 |
| *Magnification* | 81,000 | 130,000 |
| *Pixel size (Å)* | 1.10 | 0.73 |
| *Total Dose*  *(e- / Å^2^)* | 50 | 50 |
| *Exposure time (s)* | 5.72 | 4.00 |
| *Dose rate*  *(e- pixel^-1^ s^-1^)* | 10.58 | 6.65 |
| *Defocus range during data collection (μm)* | -0.5 – -3.0 | -0.5 – -3.0 |
| *Number of micrographs* | 10,005 | 7,631 |
| *Symmetry* | C1 | C1 |
| *Number of pre-refinement particles* | 1,294,603 | 204,349 |

**Table S3.** Single-particle cryo-EM model refinement parameters.

| *Sample* | M1-DD-1B2 | M1-DD-1B2 | CL-M1-TEII-1B2 |
| --- | --- | --- | --- |
| *Structure* | *Transacylation-mode* | *Elongation-mode* | *CL-M1-TEII-1B2* |
| *PDB ID* | 9PAT | 9PAV | 9PC6 |
| *EMDB ID* | EMD-71445 | EMD-71446 | EMD-71497 |
| *Resolution*  *(0.143 FSC, Å)* | 3.96 | 3.22 | 3.96 |
| *Clashscore*  *(all atoms)* | 14.91 | 10.04 | 16.76 |
| *Poor rotamers (%)* | 0.19 | 0.49 | 1.91 |
| *Ramachandran outliers (%)* | 0.00 | 0.00 | 0.00 |
| *Ramachandran favored (%)* | 96.82 | 97.11 | 97.34 |
| *MolProbity score* | 1.87 | 1.68 | 2.07 |
| *Bond length (RMSD, Å)* | 0.003 | 0.003 | 0.003 |
| *Bond angles (RMSD, °)* | 0.570 | 0.565 | 0.583 |

**Table S4.** Protein sequence key for protein sequences below.

KS = ketosynthase

AT = acyltransferase

KR = ketoreductase

DH° = inactive dehydratase

CP = carrier protein

TEII = type II thioesterase

RIFS LM (RifA)

RIFS M1 (RifA)

C-terminal Docking Domain of DEBS1 (a.k.a., EryAI) Module 2 (DD2)

N-terminal Docking Domain of DEBS2 (a.k.a., EryAII) Module 3 (DD3)

TEII (RifR)

F_ab_ 1B2

Linkers/Tags

**Protein Sequences**

*RIFS LM-M1 with C-terminal docking domain from 6-deoxyerythronolide B synthase (****LM-M1-DD****, pDC42)* **|** A-CPL-KS-AT-DH°-KR-CP1-(DD2)-His_6_

MRTDLIKPLHVALLENATRFAGKPAFADDHRTVTYGDLEARTRRLAGHLAGLGVRHGDRVAICLGNRVSTVESYFAILRAGAVGVPLNPGSATAELEHPLTDSGATVVVTDAAQAARLRLAPHVELLVTGDDVPEGAHSYDELALSEPAEPAADDLELDEPAWMFYTSGTTGRPKGVVSTQRNCLWSVASCYVPFPGLSDQDRVLWPLPLFHSLSHIACVLSATVVGASVRIADGSSADDVMRLIEAESSTFLAGVPTTYHHLVRAARQRGFSAPSLRIGLAGGAVLGAGLRSEFEETFGVPLIDAYGSTETCGAITMNPPDGARVEGSCGLAVPGVDVRVVDPDTGLDVPAGEEGEVWVSGPNVMLGYHNSPEATAAAMRDGWFRTGDLARRDDAGYFTICGRIKELIIRGGANIHPGEVEAVLRTVDGVADAAVGGVPHDTLGEVPVAYVIPGPTGFDPAALIEKCREQLSAYKVPDRILEVAHIPRTASGKIRRGLLTDEPAQLRYAATEHEEQSRHADESVAAALRARLSGLDERAQCELLEDLVRTQAADVLGQPVPDGRAFRDLGFTSLAIVELRNRLTEHTGLWLPASAVFDHPTPAALAARVRAELLGITQAVAEPVVAADPGEPIAIVGMACRLPGGVASPEDLWRLVAERVDAVSEFPGDRGWDLDSLIDPDRERAGTSYVGQGGFLHDAGEFDAGFFGISPREAVAMDPQQRLLLETSWEALENAGVDPIALKGTDTGVFSGLMGQGYGSGAVAPELEGFVTTGVASSVASGRVSYVLGLEGPAVTVDTACSSSLVAMHLAAQALRQGECSMALAGGVTVMATPGSFVEFSRQRALAPDGRCKAFAAAADGTGWSEGVGVVVLERLSVARERGHRILAVLRGSAVNQDGASNGLTAPNGLSQQRVIRRALAAAGLAPSDVDVVEAHGTGTTLGDPIEAQALLATYGQERKQPLWLGSLKSNIGHAQAAAGVAGVIKMVQALRHETLPPTLHVDKPTLEVDWSAGAIELLTEARAWPRNGRPRRAGVSSFGVSGTNAHLILEEAPAEEPVAAPELPVVPLVVSARSTESLSGQAERLASLLEGDVSLTEVAGALVSRRAVLDERAVVVAGSREEAVTGLRALNTAGSGTPGKVVWVFPGQGTQWAGMGRELLAESPVFAERIAECAAALAPWIDWSLVDVLRGEGDLGRVDVLQPACFAVMVGLAAVWESVGVRPDAVVGHSQGEIAAACVSGALSLEDAAKVVALRSQAIAAELSGRGGMASVALGEDDVVSRLVDGVEVAAVNGPSSVVIAGDAHALDATLEILSGEGIRVRRVAVDYASHTRHVEDIRDTLAETLAGISAQAPAVPFYSTVTSEWVRDAGVLDGGYWYRNLRNQVRFGAAATALLEQGHTVFVEVSAHPVTVQPLSELTGDAIGTLRREDGGLRRLLASMGELFVRGIDVDWTAMVPAAGWVDLPTYAFEHRHYWLEPAEPASAGDPLLGTVVSTPGSDRLTAVAQWSRRAQPWAVDGLVPNAALVEAAIRLGDLAGTPVVGELVVDAPVVLPRRGSREVQLIVGEPGEQRRRPIEVFSREADEPWTRHAHGTLAPAAAAVPEPAAAGDATDVTVAGLRDADRYGIHPALLDAAVRTVVGDDLLPSVWTGVSLLASGATAVTVTPTATGLRLTDPAGQPVLTVESVRGTPFVAEQGTTDALFRVDWPEIPLPTAETADFLPYEATSAEATLSALQAWLADPAETRLAVVTGDCTEPGAAAIWGLVRSAQSEHPGRIVLADLDDPAVLPAVVASGEPQVRVRNGVASVPRLTRVTPRQDARPLDPEGTVLITGGTGTLGALTARHLVTAHGVRHLVLVSRRGEAPELQEELTALGASVAIAACDVADRAQLEAVLRAIPAEHPLTAVIHTAGVLDDGVVTELTPDRLATVRRPKVDAARLLDELTREADLAAFVLFSSAAGVLGNPGQAGYAAANAELDALARQRNSLDLPAVSIAWGYWATVSGMTEHLGDADLRRNQRIGMSGLPADEGMALLDAAIATGGTLVAAKFDVAALRATAKAGGPVPPLLRGLAPLPRRAAAKTASLTERLAGLAETEQAAALLDLVRRHAAEVLGHSGAESVHSGRTFKDAGFDSLTAVELRNRLAAATGLTLSPAMIFDYPKPPALADHLRAKLFGTEVRGEAPSALAGLDALEAALPEVPATEREELVQRLERMLAALRPVAQAADASGTGANPSGDDLGEAGVDELLEALGRELDGDGNSSSVDKLAAALEHHHHHH

*RIFS LM-M1 with C-terminal TEII (a.k.a. RifR) inactivated by S2316A mutation and separated by a* (G_4_S)_8_ linker *(****LM-M1-TEII*****, pAM6)* **|** A-CPL-KS-AT-DH°-KR-CP1-(G_4_S)_8_-TEII*-His_6_

MRTDLIKPLHVALLENATRFAGKPAFADDHRTVTYGDLEARTRRLAGHLAGLGVRHGDRVAICLGNRVSTVESYFAILRAGAVGVPLNPGSATAELEHPLTDSGATVVVTDAAQAARLRLAPHVELLVTGDDVPEGAHSYDELALSEPAEPAADDLELDEPAWMFYTSGTTGRPKGVVSTQRNCLWSVASCYVPFPGLSDQDRVLWPLPLFHSLSHIACVLSATVVGASVRIADGSSADDVMRLIEAESSTFLAGVPTTYHHLVRAARQRGFSAPSLRIGLAGGAVLGAGLRSEFEETFGVPLIDAYGSTETCGAITMNPPDGARVEGSCGLAVPGVDVRVVDPDTGLDVPAGEEGEVWVSGPNVMLGYHNSPEATAAAMRDGWFRTGDLARRDDAGYFTICGRIKELIIRGGANIHPGEVEAVLRTVDGVADAAVGGVPHDTLGEVPVAYVIPGPTGFDPAALIEKCREQLSAYKVPDRILEVAHIPRTASGKIRRGLLTDEPAQLRYAATEHEEQSRHADESVAAALRARLSGLDERAQCELLEDLVRTQAADVLGQPVPDGRAFRDLGFTSLAIVELRNRLTEHTGLWLPASAVFDHPTPAALAARVRAELLGITQAVAEPVVAADPGEPIAIVGMACRLPGGVASPEDLWRLVAERVDAVSEFPGDRGWDLDSLIDPDRERAGTSYVGQGGFLHDAGEFDAGFFGISPREAVAMDPQQRLLLETSWEALENAGVDPIALKGTDTGVFSGLMGQGYGSGAVAPELEGFVTTGVASSVASGRVSYVLGLEGPAVTVDTACSSSLVAMHLAAQALRQGECSMALAGGVTVMATPGSFVEFSRQRALAPDGRCKAFAAAADGTGWSEGVGVVVLERLSVARERGHRILAVLRGSAVNQDGASNGLTAPNGLSQQRVIRRALAAAGLAPSDVDVVEAHGTGTTLGDPIEAQALLATYGQERKQPLWLGSLKSNIGHAQAAAGVAGVIKMVQALRHETLPPTLHVDKPTLEVDWSAGAIELLTEARAWPRNGRPRRAGVSSFGVSGTNAHLILEEAPAEEPVAAPELPVVPLVVSARSTESLSGQAERLASLLEGDVSLTEVAGALVSRRAVLDERAVVVAGSREEAVTGLRALNTAGSGTPGKVVWVFPGQGTQWAGMGRELLAESPVFAERIAECAAALAPWIDWSLVDVLRGEGDLGRVDVLQPACFAVMVGLAAVWESVGVRPDAVVGHSQGEIAAACVSGALSLEDAAKVVALRSQAIAAELSGRGGMASVALGEDDVVSRLVDGVEVAAVNGPSSVVIAGDAHALDATLEILSGEGIRVRRVAVDYASHTRHVEDIRDTLAETLAGISAQAPAVPFYSTVTSEWVRDAGVLDGGYWYRNLRNQVRFGAAATALLEQGHTVFVEVSAHPVTVQPLSELTGDAIGTLRREDGGLRRLLASMGELFVRGIDVDWTAMVPAAGWVDLPTYAFEHRHYWLEPAEPASAGDPLLGTVVSTPGSDRLTAVAQWSRRAQPWAVDGLVPNAALVEAAIRLGDLAGTPVVGELVVDAPVVLPRRGSREVQLIVGEPGEQRRRPIEVFSREADEPWTRHAHGTLAPAAAAVPEPAAAGDATDVTVAGLRDADRYGIHPALLDAAVRTVVGDDLLPSVWTGVSLLASGATAVTVTPTATGLRLTDPAGQPVLTVESVRGTPFVAEQGTTDALFRVDWPEIPLPTAETADFLPYEATSAEATLSALQAWLADPAETRLAVVTGDCTEPGAAAIWGLVRSAQSEHPGRIVLADLDDPAVLPAVVASGEPQVRVRNGVASVPRLTRVTPRQDARPLDPEGTVLITGGTGTLGALTARHLVTAHGVRHLVLVSRRGEAPELQEELTALGASVAIAACDVADRAQLEAVLRAIPAEHPLTAVIHTAGVLDDGVVTELTPDRLATVRRPKVDAARLLDELTREADLAAFVLFSSAAGVLGNPGQAGYAAANAELDALARQRNSLDLPAVSIAWGYWATVSGMTEHLGDADLRRNQRIGMSGLPADEGMALLDAAIATGGTLVAAKFDVAALRATAKAGGPVPPLLRGLAPLPRRAAAKTASLTERLAGLAETEQAAALLDLVRRHAAEVLGHSGAESVHSGRTFKDAGFDSLTAVELRNRLAAATGLTLSPAMIFDYPKPPALADHLRAKLFGSAASGGGGSGGGGSGGGGSGGGGSGGGGSGGGGSGGGGSGGGGSHRPEAEKWLRRFERAPDARARLVCLPHAGGSASFFFPLAKALAPAVEVLAVQYPGRQDRRHEPPVDSIGGLTNRLLEVLRPFGDRPLALFGHAMGAIIGYELALRMPEAGLPAPVHLFASGRRAPSRYRDDDVRGASDERLVAELRKLGGSDAAMLADPELLAMVLPAIRSDYRAVETYRHEPGRRVDCPVTVFTGDHDPRVSVGEARAWEEHTTGPADLRVLPGGHFFLVDQAAPMIATMTEKLAGPALTGSTGGNSGNSSSVDKLAAALEHHHHHH

*RIFS M1 with N- and C-terminal docking domains from 6-deoxyerythronolide B synthase (****M1-DD****, pDC43)* **|** (DD3)-KS-AT-DH°-KR-CP1-(DD2)-His_6_

MASTDSEKVAEYLRRATLDLRAARQRIRELEGEPIAIVGMACRLPGGVASPEDLWRLVAERVDAVSEFPGDRGWDLDSLIDPDRERAGTSYVGQGGFLHDAGEFDAGFFGISPREAVAMDPQQRLLLETSWEALENAGVDPIALKGTDTGVFSGLMGQGYGSGAVAPELEGFVTTGVASSVASGRVSYVLGLEGPAVTVDTACSSSLVAMHLAAQALRQGECSMALAGGVTVMATPGSFVEFSRQRALAPDGRCKAFAAAADGTGWSEGVGVVVLERLSVARERGHRILAVLRGSAVNQDGASNGLTAPNGLSQQRVIRRALAAAGLAPSDVDVVEAHGTGTTLGDPIEAQALLATYGQERKQPLWLGSLKSNIGHAQAAAGVAGVIKMVQALRHETLPPTLHVDKPTLEVDWSAGAIELLTEARAWPRNGRPRRAGVSSFGVSGTNAHLILEEAPAEEPVAAPELPVVPLVVSARSTESLSGQAERLASLLEGDVSLTEVAGALVSRRAVLDERAVVVAGSREEAVTGLRALNTAGSGTPGKVVWVFPGQGTQWAGMGRELLAESPVFAERIAECAAALAPWIDWSLVDVLRGEGDLGRVDVLQPACFAVMVGLAAVWESVGVRPDAVVGHSQGEIAAACVSGALSLEDAAKVVALRSQAIAAELSGRGGMASVALGEDDVVSRLVDGVEVAAVNGPSSVVIAGDAHALDATLEILSGEGIRVRRVAVDYASHTRHVEDIRDTLAETLAGISAQAPAVPFYSTVTSEWVRDAGVLDGGYWYRNLRNQVRFGAAATALLEQGHTVFVEVSAHPVTVQPLSELTGDAIGTLRREDGGLRRLLASMGELFVRGIDVDWTAMVPAAGWVDLPTYAFEHRHYWLEPAEPASAGDPLLGTVVSTPGSDRLTAVAQWSRRAQPWAVDGLVPNAALVEAAIRLGDLAGTPVVGELVVDAPVVLPRRGSREVQLIVGEPGEQRRRPIEVFSREADEPWTRHAHGTLAPAAAAVPEPAAAGDATDVTVAGLRDADRYGIHPALLDAAVRTVVGDDLLPSVWTGVSLLASGATAVTVTPTATGLRLTDPAGQPVLTVESVRGTPFVAEQGTTDALFRVDWPEIPLPTAETADFLPYEATSAEATLSALQAWLADPAETRLAVVTGDCTEPGAAAIWGLVRSAQSEHPGRIVLADLDDPAVLPAVVASGEPQVRVRNGVASVPRLTRVTPRQDARPLDPEGTVLITGGTGTLGALTARHLVTAHGVRHLVLVSRRGEAPELQEELTALGASVAIAACDVADRAQLEAVLRAIPAEHPLTAVIHTAGVLDDGVVTELTPDRLATVRRPKVDAARLLDELTREADLAAFVLFSSAAGVLGNPGQAGYAAANAELDALARQRNSLDLPAVSIAWGYWATVSGMTEHLGDADLRRNQRIGMSGLPADEGMALLDAAIATGGTLVAAKFDVAALRATAKAGGPVPPLLRGLAPLPRRAAAKTASLTERLAGLAETEQAAALLDLVRRHAAEVLGHSGAESVHSGRTFKDAGFDSLTAVELRNRLAAATGLTLSPAMIFDYPKPPALADHLRAKLFGTEVRGEAPSALAGLDALEAALPEVPATEREELVQRLERMLAALRPVAQAADASGTGANPSGDDLGEAGVDELLEALGRELDGDGNSSSVDKLAAALEHHHHHH

*RIFS M1 with N-terminal docking domain from 6-deoxyerythronolide B synthase and C-terminal TEII (a.k.a. RifR) separated by a* (G_4_S)_2_ linker *(****M1-TEII****, pDC85)* **|** (DD3)-KS-AT-DH°-KR-CP1-(G_4_S)_2_-TEII-His_6_

MASTDSEKVAEYLRRATLDLRAARQRIRELEGEPIAIVGMACRLPGGVASPEDLWRLVAERVDAVSEFPGDRGWDLDSLIDPDRERAGTSYVGQGGFLHDAGEFDAGFFGISPREAVAMDPQQRLLLETSWEALENAGVDPIALKGTDTGVFSGLMGQGYGSGAVAPELEGFVTTGVASSVASGRVSYVLGLEGPAVTVDTACSSSLVAMHLAAQALRQGECSMALAGGVTVMATPGSFVEFSRQRALAPDGRCKAFAAAADGTGWSEGVGVVVLERLSVARERGHRILAVLRGSAVNQDGASNGLTAPNGLSQQRVIRRALAAAGLAPSDVDVVEAHGTGTTLGDPIEAQALLATYGQERKQPLWLGSLKSNIGHAQAAAGVAGVIKMVQALRHETLPPTLHVDKPTLEVDWSAGAIELLTEARAWPRNGRPRRAGVSSFGVSGTNAHLILEEAPAEEPVAAPELPVVPLVVSARSTESLSGQAERLASLLEGDVSLTEVAGALVSRRAVLDERAVVVAGSREEAVTGLRALNTAGSGTPGKVVWVFPGQGTQWAGMGRELLAESPVFAERIAECAAALAPWIDWSLVDVLRGEGDLGRVDVLQPACFAVMVGLAAVWESVGVRPDAVVGHSQGEIAAACVSGALSLEDAAKVVALRSQAIAAELSGRGGMASVALGEDDVVSRLVDGVEVAAVNGPSSVVIAGDAHALDATLEILSGEGIRVRRVAVDYASHTRHVEDIRDTLAETLAGISAQAPAVPFYSTVTSEWVRDAGVLDGGYWYRNLRNQVRFGAAATALLEQGHTVFVEVSAHPVTVQPLSELTGDAIGTLRREDGGLRRLLASMGELFVRGIDVDWTAMVPAAGWVDLPTYAFEHRHYWLEPAEPASAGDPLLGTVVSTPGSDRLTAVAQWSRRAQPWAVDGLVPNAALVEAAIRLGDLAGTPVVGELVVDAPVVLPRRGSREVQLIVGEPGEQRRRPIEVFSREADEPWTRHAHGTLAPAAAAVPEPAAAGDATDVTVAGLRDADRYGIHPALLDAAVRTVVGDDLLPSVWTGVSLLASGATAVTVTPTATGLRLTDPAGQPVLTVESVRGTPFVAEQGTTDALFRVDWPEIPLPTAETADFLPYEATSAEATLSALQAWLADPAETRLAVVTGDCTEPGAAAIWGLVRSAQSEHPGRIVLADLDDPAVLPAVVASGEPQVRVRNGVASVPRLTRVTPRQDARPLDPEGTVLITGGTGTLGALTARHLVTAHGVRHLVLVSRRGEAPELQEELTALGASVAIAACDVADRAQLEAVLRAIPAEHPLTAVIHTAGVLDDGVVTELTPDRLATVRRPKVDAARLLDELTREADLAAFVLFSSAAGVLGNPGQAGYAAANAELDALARQRNSLDLPAVSIAWGYWATVSGMTEHLGDADLRRNQRIGMSGLPADEGMALLDAAIATGGTLVAAKFDVAALRATAKAGGPVPPLLRGLAPLPRRAAAKTASLTERLAGLAETEQAAALLDLVRRHAAEVLGHSGAESVHSGRTFKDAGFDSLTAVELRNRLAAATGLTLSPAMIFDYPKPPALADHLRAKLFGSGGGGSGGGGSHRPEAEKWLRRFERAPDARARLVCLPHAGGSASFFFPLAKALAPAVEVLAVQYPGRQDRRHEPPVDSIGGLTNRLLEVLRPFGDRPLALFGHSMGAIIGYELALRMPEAGLPAPVHLFASGRRAPSRYRDDDVRGASDERLVAELRKLGGSDAAMLADPELLAMVLPAIRSDYRAVETYRHEPGRRVDCPVTVFTGDHDPRVSVGEARAWEEHTTGPADLRVLPGGHFFLVDQAAPMIATMTEKLAGPALTGSTGGNSGNSSSVDKLAAALEHHHHHH

*RIFS M1 with N-terminal docking domain from 6-deoxyerythronolide B synthase and C-terminal TEII (a.k.a. RifR) inactivated by S2316A mutation and separated by a* (G_4_S)_8_ linker *(****M1-TEII*****, pDC147)* **|** (DD3)-KS-AT-DH°-KR-CP1-(G_4_S)_8_-TEII*-His_6_

MASTDSEKVAEYLRRATLDLRAARQRIRELEGEPIAIVGMACRLPGGVASPEDLWRLVAERVDAVSEFPGDRGWDLDSLIDPDRERAGTSYVGQGGFLHDAGEFDAGFFGISPREAVAMDPQQRLLLETSWEALENAGVDPIALKGTDTGVFSGLMGQGYGSGAVAPELEGFVTTGVASSVASGRVSYVLGLEGPAVTVDTACSSSLVAMHLAAQALRQGECSMALAGGVTVMATPGSFVEFSRQRALAPDGRCKAFAAAADGTGWSEGVGVVVLERLSVARERGHRILAVLRGSAVNQDGASNGLTAPNGLSQQRVIRRALAAAGLAPSDVDVVEAHGTGTTLGDPIEAQALLATYGQERKQPLWLGSLKSNIGHAQAAAGVAGVIKMVQALRHETLPPTLHVDKPTLEVDWSAGAIELLTEARAWPRNGRPRRAGVSSFGVSGTNAHLILEEAPAEEPVAAPELPVVPLVVSARSTESLSGQAERLASLLEGDVSLTEVAGALVSRRAVLDERAVVVAGSREEAVTGLRALNTAGSGTPGKVVWVFPGQGTQWAGMGRELLAESPVFAERIAECAAALAPWIDWSLVDVLRGEGDLGRVDVLQPACFAVMVGLAAVWESVGVRPDAVVGHSQGEIAAACVSGALSLEDAAKVVALRSQAIAAELSGRGGMASVALGEDDVVSRLVDGVEVAAVNGPSSVVIAGDAHALDATLEILSGEGIRVRRVAVDYASHTRHVEDIRDTLAETLAGISAQAPAVPFYSTVTSEWVRDAGVLDGGYWYRNLRNQVRFGAAATALLEQGHTVFVEVSAHPVTVQPLSELTGDAIGTLRREDGGLRRLLASMGELFVRGIDVDWTAMVPAAGWVDLPTYAFEHRHYWLEPAEPASAGDPLLGTVVSTPGSDRLTAVAQWSRRAQPWAVDGLVPNAALVEAAIRLGDLAGTPVVGELVVDAPVVLPRRGSREVQLIVGEPGEQRRRPIEVFSREADEPWTRHAHGTLAPAAAAVPEPAAAGDATDVTVAGLRDADRYGIHPALLDAAVRTVVGDDLLPSVWTGVSLLASGATAVTVTPTATGLRLTDPAGQPVLTVESVRGTPFVAEQGTTDALFRVDWPEIPLPTAETADFLPYEATSAEATLSALQAWLADPAETRLAVVTGDCTEPGAAAIWGLVRSAQSEHPGRIVLADLDDPAVLPAVVASGEPQVRVRNGVASVPRLTRVTPRQDARPLDPEGTVLITGGTGTLGALTARHLVTAHGVRHLVLVSRRGEAPELQEELTALGASVAIAACDVADRAQLEAVLRAIPAEHPLTAVIHTAGVLDDGVVTELTPDRLATVRRPKVDAARLLDELTREADLAAFVLFSSAAGVLGNPGQAGYAAANAELDALARQRNSLDLPAVSIAWGYWATVSGMTEHLGDADLRRNQRIGMSGLPADEGMALLDAAIATGGTLVAAKFDVAALRATAKAGGPVPPLLRGLAPLPRRAAAKTASLTERLAGLAETEQAAALLDLVRRHAAEVLGHSGAESVHSGRTFKDAGFDSLTAVELRNRLAAATGLTLSPAMIFDYPKPPALADHLRAKLFGSAASGGGGSGGGGSGGGGSGGGGSGGGGSGGGGSGGGGSGGGGSHRPEAEKWLRRFERAPDARARLVCLPHAGGSASFFFPLAKALAPAVEVLAVQYPGRQDRRHEPPVDSIGGLTNRLLEVLRPFGDRPLALFGHAMGAIIGYELALRMPEAGLPAPVHLFASGRRAPSRYRDDDVRGASDERLVAELRKLGGSDAAMLADPELLAMVLPAIRSDYRAVETYRHEPGRRVDCPVTVFTGDHDPRVSVGEARAWEEHTTGPADLRVLPGGHFFLVDQAAPMIATMTEKLAGPALTGSTGGNSGNSSSVDKLAAALEHHHHHH

*RIFS M1 with N- and C-terminal docking domains from the 6-deoxyerythronolide B synthase and a 38-residue deletion (Δ1588–1625) within the C-terminal docking domain corresponding to its dimeric α-helical fragment (****M1-ΔDD****, pDC155)* **|** (DD3)-KS-AT-DH°-KR-CP1-(ΔDD2)-His_6_

MASTDSEKVAEYLRRATLDLRAARQRIRELEGEPIAIVGMACRLPGGVASPEDLWRLVAERVDAVSEFPGDRGWDLDSLIDPDRERAGTSYVGQGGFLHDAGEFDAGFFGISPREAVAMDPQQRLLLETSWEALENAGVDPIALKGTDTGVFSGLMGQGYGSGAVAPELEGFVTTGVASSVASGRVSYVLGLEGPAVTVDTACSSSLVAMHLAAQALRQGECSMALAGGVTVMATPGSFVEFSRQRALAPDGRCKAFAAAADGTGWSEGVGVVVLERLSVARERGHRILAVLRGSAVNQDGASNGLTAPNGLSQQRVIRRALAAAGLAPSDVDVVEAHGTGTTLGDPIEAQALLATYGQERKQPLWLGSLKSNIGHAQAAAGVAGVIKMVQALRHETLPPTLHVDKPTLEVDWSAGAIELLTEARAWPRNGRPRRAGVSSFGVSGTNAHLILEEAPAEEPVAAPELPVVPLVVSARSTESLSGQAERLASLLEGDVSLTEVAGALVSRRAVLDERAVVVAGSREEAVTGLRALNTAGSGTPGKVVWVFPGQGTQWAGMGRELLAESPVFAERIAECAAALAPWIDWSLVDVLRGEGDLGRVDVLQPACFAVMVGLAAVWESVGVRPDAVVGHSQGEIAAACVSGALSLEDAAKVVALRSQAIAAELSGRGGMASVALGEDDVVSRLVDGVEVAAVNGPSSVVIAGDAHALDATLEILSGEGIRVRRVAVDYASHTRHVEDIRDTLAETLAGISAQAPAVPFYSTVTSEWVRDAGVLDGGYWYRNLRNQVRFGAAATALLEQGHTVFVEVSAHPVTVQPLSELTGDAIGTLRREDGGLRRLLASMGELFVRGIDVDWTAMVPAAGWVDLPTYAFEHRHYWLEPAEPASAGDPLLGTVVSTPGSDRLTAVAQWSRRAQPWAVDGLVPNAALVEAAIRLGDLAGTPVVGELVVDAPVVLPRRGSREVQLIVGEPGEQRRRPIEVFSREADEPWTRHAHGTLAPAAAAVPEPAAAGDATDVTVAGLRDADRYGIHPALLDAAVRTVVGDDLLPSVWTGVSLLASGATAVTVTPTATGLRLTDPAGQPVLTVESVRGTPFVAEQGTTDALFRVDWPEIPLPTAETADFLPYEATSAEATLSALQAWLADPAETRLAVVTGDCTEPGAAAIWGLVRSAQSEHPGRIVLADLDDPAVLPAVVASGEPQVRVRNGVASVPRLTRVTPRQDARPLDPEGTVLITGGTGTLGALTARHLVTAHGVRHLVLVSRRGEAPELQEELTALGASVAIAACDVADRAQLEAVLRAIPAEHPLTAVIHTAGVLDDGVVTELTPDRLATVRRPKVDAARLLDELTREADLAAFVLFSSAAGVLGNPGQAGYAAANAELDALARQRNSLDLPAVSIAWGYWATVSGMTEHLGDADLRRNQRIGMSGLPADEGMALLDAAIATGGTLVAAKFDVAALRATAKAGGPVPPLLRGLAPLPRRAAAKTASLTERLAGLAETEQAAALLDLVRRHAAEVLGHSGAESVHSGRTFKDAGFDSLTAVELRNRLAAATGLTLSPAMIFDYPKPPALADHLRAKLFGTEVRGEAVAQAADASGTGANPSGDDLGEAGVDELLEALGRELDGDGNSSSVDKLAAALEHHHHHH

*F_ab_ 1B2 heavy chain with a C-terminal His_6_ and FLAG tag*

MAEVQLVQSGGGLVQPGRSLRLSCTASGFTFGDYAMSWVRQAPGKGLEWVGFIRSKAYGGTTEYAASVKGRFTISRDDSKSIAYLQMNSLKTEDTAVYYCTRGGTLFDYWGQGTLVTVSSASTKGPSVFPLAPSSKSTSGGTAALGCLVKDYFPEPVTVSWNSGALTSGVHTFPAVLQSSGLYSLSSVVTVPSSSLGTQTYICNVNHKPSNTKVDKKVEPKSCAALVPRGSAHHHHHHAADYKDDDDKA

*F_ab_ 1B2 light chain*

LFAIPLVVPFYSHSALDVVMTQSPLSLPVTPGEPASISCRSSQSLLHSNGYNYLDWYLQKPGQSPQLLIYLGSNRASGVPDRFSGSGSGTDFTLKISRVEAEDVGVYYCMQSLQTPRLTFGPGTKVDIKRTVAAPSVFIFPPSDEQLKSGTASVVCLLNNFYPRGAKVQWKVDNALQSGNSQESVTEQDSKDSTYSLSSTLTLSKADYEKHKVYACEVTHQGLSSPVTKSFNRGEC
